# Follistatin improves vascular function by inhibiting oxidative stress and inducing browning of perivascular adipose tissue in essential hypertension

**DOI:** 10.64898/2025.12.31.697255

**Authors:** Ann Kuganathan, Vincent Lu, Melissa MacDonald, Sofia Farkona, Tracey Campbell, Madoka Akimoto, Bo Gao, Kieran Manion, Ana Konvalinka, Jeffrey Dickhout, Joan C. Krepinsky

**Author notes:** **Correspondence to:** Joan C. Krepinsky St. Joseph’s Hospital 50 Charlton Ave East, T3311 Hamilton, ON L8N 4A6, Canada. Contributed equally to this work (Jeffrey Dickhout, Joan C. Krepinsky).

## Abstract

**Background:** Essential hypertension, characterized by vascular dysfunction, remains a leading modifiable cause of death globally. Perivascular adipose tissue (PVAT), which normally reduces vasoconstriction, becomes less anticontractile in hypertension due at least in part to increased reactive oxygen species (ROS). Brown PVAT has emerged as a protective regulator of vascular tone. Follistatin, an activin antagonist, induces browning of peripheral adipose tissue depots via AMPK. We recently reported improved blood pressure, vascular function and reduced vascular ROS by follistatin in the spontaneously hypertensive rat (SHR) model of essential hypertension. Here, we investigate whether follistatin reduces ROS and induces browning in SHR PVAT to restore PVAT function.

**Methods:** SHR were treated with vehicle, follistatin or hydralazine for 8 weeks. Mesenteric white and thoracic brown PVAT from SHR and normotensive Wistar-Kyoto were utilized, with wire myography used to assess vascular function. Nitric oxide (NO) and nitrite were measured in PVAT using DAF-2 FM fluorescence or Griess reagent, respectively. PVAT ROS and browning markers were assessed biochemically and via immunohistochemistry. PVAT was treated ex vivo for 3 days to assess mechanisms of browning. Unbiased proteomic analysis of PVAT was performed using liquid chromatography-mass spectrometry.

**Results:** SHR PVAT dysfunction, manifest as a pro-contractile effect, was inhibited by follistatin through reducing ROS, enhancing NO bioavailability and inducing browning. Activin neutralization and AMPK phosphorylation mediated the beneficial effect of follistatin on SHR PVAT. Hydralazine-induced blood pressure reduction did not replicate follistatin effects, suggesting vascular benefits of follistatin are mediated by direct PVAT modulation. Proteomic analysis showed that follistatin shifts PVAT proteome towards normal state, upregulating processes associated with adipose tissue browning.

**Conclusion:** Follistatin restores PVAT-mediated vascular relaxation via ROS reduction, activin neutralization and AMPK-dependent browning, positioning the potential for PVAT as a therapeutic target for vascular dysfunction in essential hypertension.

## Introduction

Hypertension is a leading modifiable risk factor for cardiovascular morbidity and mortality worldwide^1^, however, current antihypertensive therapies often fail to achieve optimal blood pressure (BP) control^2^. Despite its prevalence, the underlying cause of hypertension remains unknown in most cases, termed essential hypertension^3^. While characterized by vascular dysfunction^3^, the important contribution of perivascular adipose tissue (PVAT) is becoming increasingly recognized. PVAT surrounds most blood vessels, exerting a paracrine effect through the release of vasoactive substances. In normal conditions, PVAT exerts an anticontractile effect. In hypertension, PVAT exhibits increased oxidative stress resulting in augmented release of contractile factors^4^. Notably, PVAT dysfunction precedes the development of hypertension in the spontaneously hypertensive rat (SHR) model of essential hypertension, suggesting a potential role in its pathophysiology^5^.

Adipose tissue is heterogenous and is broadly classified as either white, brown or beige. White adipose tissue (WAT) accumulates in metabolic diseases such as obesity and is characterized by excessive oxidative stress and inflammation^6^. In contrast, brown adipose tissue (BAT) has a more potent anticontractile effect and protects against obesity-induced inflammation and vascular dysfunction^7^. In individuals, the presence of BAT is correlated with a lower prevalence of hypertension^8^. Upon β_3_-adrenergic receptor activation or cold exposure, WAT undergoes a phenotypic transformation into BAT, a process referred to as adipose tissue browning. PVAT browning has been associated with BP reduction^9^.

We recently found that the glycoprotein follistatin significantly lowers BP and improves vessel function via oxidative stress inhibition in essential and secondary hypertension models^10^. Follistatin inhibits several members of the TGFβ superfamily with potent effects against activins A and B^11^. Activins are regulators of oxidative stress and vessel contraction which have been implicated in the pathophysiology of hypertension^12,13,14^. Activin A has also been shown to suppress differentiation of brown adipocytes^15^. Of relevance, follistatin induces the differentiation of brown adipocytes and the browning of peripheral WAT in vitro^16^ and in vivo^17,18^ respectively via AMP-activated protein kinase (AMPK). Whether follistatin induces PVAT browning is unknown.

In this study, we investigated the effects of follistatin on PVAT structure and function in the SHR model of essential hypertension. We demonstrate that follistatin significantly improves PVAT-mediated vascular function by reducing reactive oxygen species (ROS) and promoting PVAT browning via activin neutralization and AMPK phosphorylation. These effects are not replicated by BP lowering alone with hydralazine. Together, these findings highlight a novel mechanism by which follistatin restores vascular homeostasis and support the therapeutic potential of targeting PVAT in hypertension.

## Methods

Detailed Methods are described in Supplemental Material.

### Animal studies

All experiments adhered to guidelines set by McMaster University, the Canadian Council on Animal Care (CCAC) and ARRIVE and were approved by the McMaster University Animals for Research Ethics Board.

### General statistical analysis

Statistical analyses were conducted using GraphPad Prism10. To assess statistical significance for BP and histology data, a one-way ANOVA was employed. Weekly BP data were analyzed using a repeated measures test. Myograph data were analyzed using a non-linear regression model. Two-way ANOVA between two groups was used to compare difference between curves. Imaging data were evaluated using one-way ANOVA with a post hoc Tukey correction for multiple comparisons. For comparisons between two groups, a Student’s t-test was utilized. Proteomic data was analyzed using Proteome Discoverer using the Rattus norvegicus fasta database. Gene Ontology (GO) for biological processes was performed on differentially expressed proteins using Database for Annotation Visualization and Integrated Discovery (DAVID). A p-value of less than 0.05 was considered statistically significant. Data are presented as mean ± SEM, with the number of samples (n) indicated in figure captions or as individual data points in the graphs.

## Results

### Follistatin inhibits the procontractile effect of SHR PVAT

We first determined the effect of PVAT on smooth muscle contraction to KCl in first-order mesenteric arteries isolated from SHR rats treated for 8 weeks with vehicle or follistatin, or normotensive rats treated for 8 weeks with vehicle. PVAT exhibited a significant anticontractile effect on WKY vessels (**Figure 1A, B)**. In contrast, SHR PVAT elicited a marked procontractile response which was reversed by follistatin **(Figure 1A-C)**. The lack of an anticontractile effect by SHR PVAT was also seen in response to the “_1_ adrenergic receptor agonist phenylephrine (**Figure 1D)**, which was partially restored by follistatin **(Figure 1E, F)**. To eliminate the vasoactive contribution of the endothelium, we assessed KCl response of endothelium-denuded SHR vessels with or without PVAT. Endothelium removal in SHR vessels augmented contraction to KCl **(Figure 1G)** suggesting that endothelium-derived vasodilatory factors modulate vascular tone in the SHR. Notably, the procontractile effect of denuding the endothelium was further exacerbated in the presence of SHR PVAT, with PVAT effects reversed by follistatin **(Figure 1H)**. These data support beneficial effects of follistatin on restoring normal PVAT regulation of vascular function.

**Figure 1.**
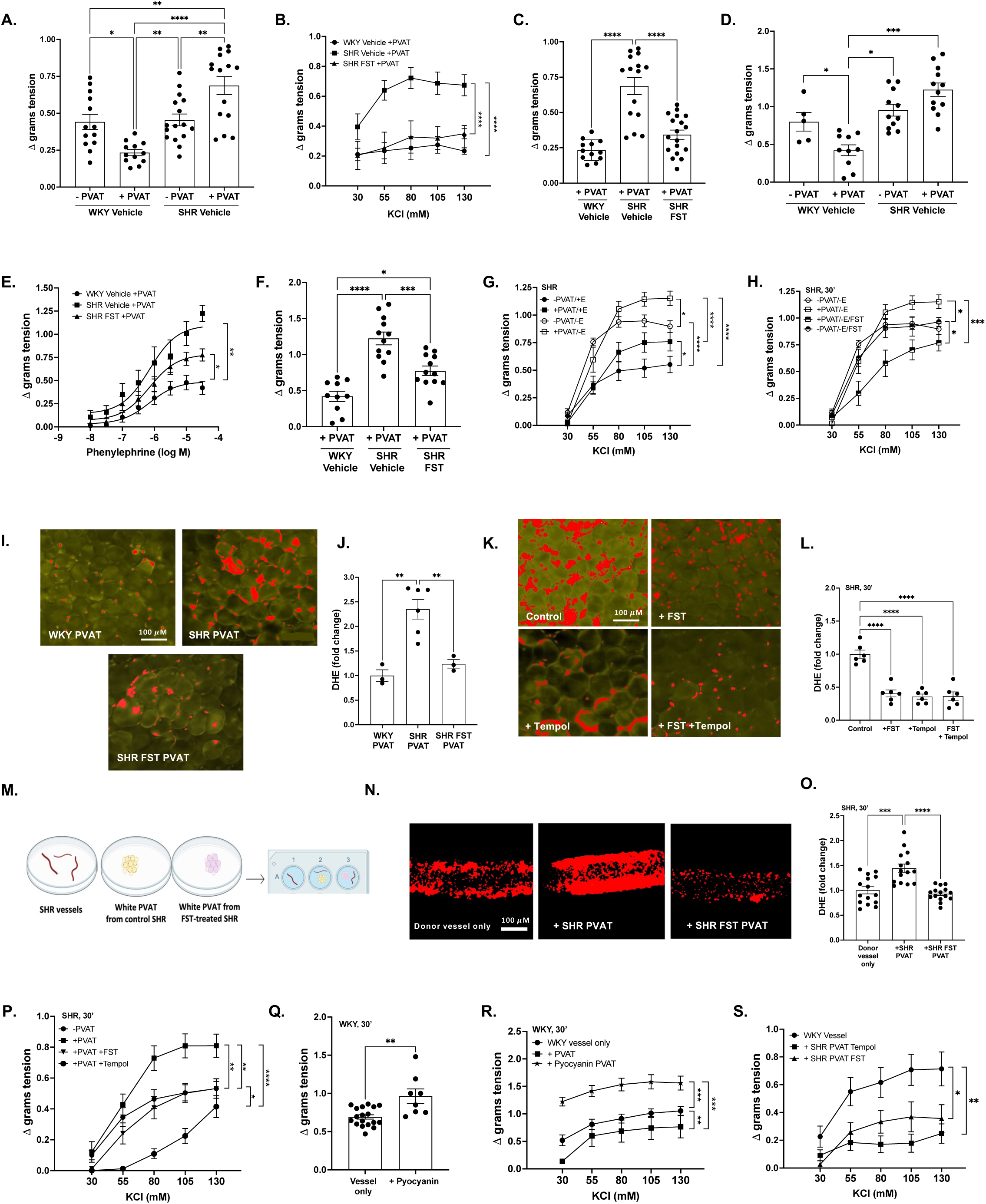
The procontractile effect of SHR PVAT, associated with increased oxidative stress, is inhibited by follistatin. **A.** Response to maximal KCl dose (125 mM) in first-order mesenteric vessels with or without PVAT from vehicle-treated WKY or SHR. **B.** KCl dose response curves (30 – 130 mM) in vessels with PVAT from treated rats, with maximal response quantified in **C. D.** Response to maximal phenylephrine dose (3×10^-5^ M) in vessels with or without PVAT from treated rats. **E.** Phenylephrine dose response curves (10^-8^ - 3×10^-5^ M) in vessels with PVAT from treated rats with maximal response quantified in **F. G.** KCl dose response curves (30 – 130 mM) in SHR vessels with either PVAT, endothelium or both removed. **H.** KCl dose response curves (30 – 130 mM) in endothelium-denuded SHR vessels treated acutely with or without follistatin for 30 minutes. **I.** Representative fluorescence images of mesenteric PVAT from treated rats stained with dihydroethidium (DHE), quantified in **J. K.** Representative fluorescence images of mesenteric PVAT treated with either follistatin, Tempol or both and stained with DHE, quantified in **L. M.** Schematic of coincubation treatment setup. **N.** Representative fluorescence images of first-order mesenteric SHR donor vessels incubated for 30 minutes with mesenteric PVAT from either vehicle- or follistatin-treated SHRs, stained with DHE and quantified in **O. P.** KCl dose response curves (30 – 130 mM) in SHR vessels with or without PVAT pretreated with either follistatin or Tempol for 30 minutes. **Q.** Response to maximal KCl dose (125 mM) in WKY vessels treated with HBSS or pyocyanin for 30 minutes. **R.** KCl dose response curves (30 – 130 mM) in WKY vessels incubated with mesenteric WKY PVAT pretreated for 30 minutes with HBSS or pyocyanin. **S.** KCl dose response curves (30 – 130 mM) in WKY vessels incubated with mesenteric SHR PVAT pretreated for 30 minutes with follistatin or Tempol. Contractility values are from 5-10 rats per group, 1-3 vessels per rat. *\*p*<0.05, *\*\*p* <0.01, *\*\*\*p* <0.001, *\*\*\*\*p* <0.0001.

### PVAT ROS induces vascular oxidative stress and vasoconstriction

We previously showed that follistatin inhibits ROS in SHR vessels^10^. We thus sought to determine whether a similar effect would be observed in PVAT. DHE fluorescence imaging showed increased ROS in SHR PVAT, which was inhibited to control levels by follistatin **(Figure 1I, J)**. Follistatin also decreased proinflammatory markers CD68 and iNOS **(Supplementary Figure 1A-D)**, and increased CD206, an anti-inflammatory marker in SHR PVAT **(Supplementary Figure 1E, F)**. We next investigated the effects of short-term incubation of follistatin on ROS levels in ex vivo SHR PVAT. As shown in **Figure 1K, L**, follistatin and the ROS inhibitor Tempol both reduced DHE fluorescence to similar levels, without an additive effect. To determine whether PVAT ROS could induce oxidative stress in the associated vessel, we performed PVAT transfer experiments. PVAT obtained from an SHR treated for 8 weeks with vehicle or follistatin were incubated with a donor vessel obtained from an untreated SHR to remove the confounding effect of follistatin on the vessel itself **(Figure 1M)**. After 30 minutes of coincubation, PVAT from vehicle-treated SHR increased vessel oxidative stress, measured by DHE. This was not seen in PVAT from follistatin-treated SHR **(Figure 1N, O)**. These findings suggest that PVAT ROS can induce vascular oxidative stress in this model.

To determine whether inhibiting ROS resulted in reduced vessel contraction, we treated SHR vessels harvested with intact PVAT with Tempol or follistatin for 30 minutes prior to myography studies. As seen in **Figure 1P**, both Tempol and follistatin reduced the procontractile effect of PVAT, although Tempol reductions were greater. These data suggest that oxidative stress can increase vessel constriction. Indeed, exposure of WKY vessels to the ROS inducer pyocyanin, increased constriction to KCl **(Figure 1Q)**. Furthermore, isolated PVAT from WKY vessels pretreated with pyocyanin augmented WKY vessel contraction to KCl **(Figure 1R)**. Conversely, the treatment of detached SHR PVAT with either follistatin or Tempol restored its anticontractile effect when co-incubated with WKY vessels stimulated with KCl **(Figure 1S)**. These findings show the potential of PVAT ROS to directly influence vascular tone and suggest that the augmented vessel contraction induced by SHR PVAT is driven at least in part by oxidative stress that is reversible by follistatin.

### Activin upregulation contributes to increased ROS in SHR PVAT

The follistatin-288 isoform used in this study is likely to deposit in tissue through its affinity for heparan sulfate proteoglycans^19^. Follistatin is also endogenously produced. We thus characterized the abundance of follistatin as well as activins in PVAT by immunohistochemistry. While follistatin was significantly reduced in SHR compared to WKY PVAT, follistatin administration increased SHR PVAT follistatin levels to greater than those seen in WKY **(Figure 2A, B)**. Concordantly, the increased expression of activins A and B in SHR PVAT was reduced by follistatin **(Figure 2C-F)**.

**Figure 2:**
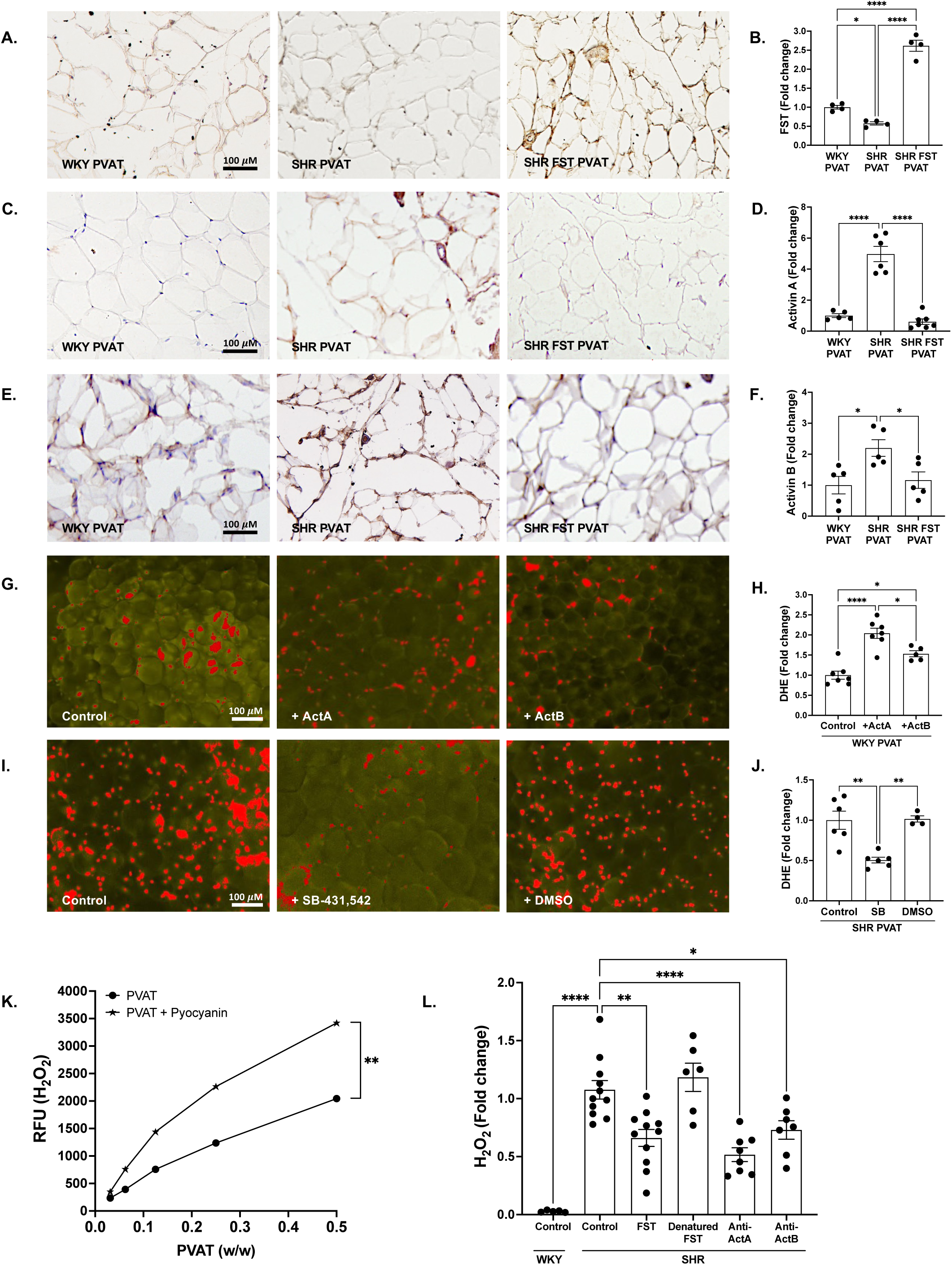
Follistatin inhibits oxidative stress in PVAT by neutralizing activins. Representative cross-sectional images of mesenteric PVAT from chronically treated rats stained for **A.** follistatin, **C.** activin A, **E.** activin B, quantified in accompanying graphs **B., D.**, and **F. G.** Representative fluorescence images of WKY mesenteric PVAT treated acutely with activin A or B for 30 minutes, quantified in **H. I.** Representative fluorescence images of SHR mesenteric PVAT treated acutely with SB-431,542 or DMSO for 30 minutes, quantified in **J. K.** Fold change in H_2_O_2_ levels assessed using Amplex Red in WKY mesenteric PVAT treated for 30 minutes with HBSS or pyocyanin. **L.** Fold change in H_2_O_2_ levels measured in mesenteric PVAT from WKY or SHR treated for 30 minutes with either HBSS, follistatin, heat-denatured follistatin, or neutralizing antibodies to activin A or activin B. *\*p*<0.05, *\*\*p* <0.01, *\*\*\*\*p* <0.0001.

We previously demonstrated increased vascular ROS upon acute treatment (30 min) with activin A^10^. To determine whether activins could induce ROS in PVAT, we treated WKY PVAT with activin A or B. As shown in **Figure 2G, H**, both increased ROS as measured by DHE. Furthermore, in SHR PVAT the activin receptor blocker SB-431,542 lowered ROS, demonstrating contribution of activins to PVAT oxidative stress in the SHR **(Figure 2I, J)**.

Vascular ROS are primarily produced by NADPH oxidases (NOX)^20^. Of the various NOX isoforms, NOX4 is abundantly expressed in all parts of the vessels, primarily producing hydrogen peroxide (H_2_O_2_)^20^. We thus sought to identify the contribution of H_2_O_2_ to the DHE signal using Amplex Red. We first confirmed an increase in fluorescence in WKY PVAT treated with pyocyanin, indicative of increased H_2_O_2_ production **(Figure 2K)**. SHR PVAT were found to have elevated H_2_O_2_ levels compared to WKY PVAT, and these were reduced by follistatin, but not by heat denatured follistatin **(Figure 2L)**. These data indicate that the tertiary structure of follistatin is required for its inhibition of ROS. Activin A or B neutralization also significantly reduced H_2_O_2_ levels in SHR PVAT, supporting their role as important mediators of ROS production in SHR PVAT.

### Follistatin increases nitric oxide (NO) bioavailability in SHR PVAT

Superoxide generation by NADPH oxidase, by reacting with and reducing NO bioavailability, is also an important contributor to increased vasoconstriction in hypertension^21,22^. Superoxide reaction with NO generates peroxynitrite which can then react with tyrosine residues in proteins to form nitrotyrosine. Immunohistochemistry detection of nitrotyrosine was significantly increased in SHR compared to WKY PVAT, and this was reduced by follistatin **(Figure 3A, B)**. Immunohistochemistry further showed that follistatin increased expression of the primary vascular NO producer eNOS **(Figure 3C, D)**. To determine if follistatin altered NO bioavailability, we used the DAF-2 FM fluorescence probe to directly detect NO. Upon addition of the NO donor, NONOate, SHR PVAT showed significantly reduced fluorescence intensity compared with WKY PVAT. This indicates that NO is rapidly taken up in SHR PVAT, likely by ROS, thus reducing its bioavailability **(Figure 3E)**. Pretreatment of SHR PVAT with follistatin for 30 minutes increased fluorescence, indicating an increase in NO bioavailability. Tempol restored NO bioavailability to normal levels in SHR PVAT, supporting a key role for oxidative stress in its reduction **(Figure 3E, F)**.

**Figure 3:**
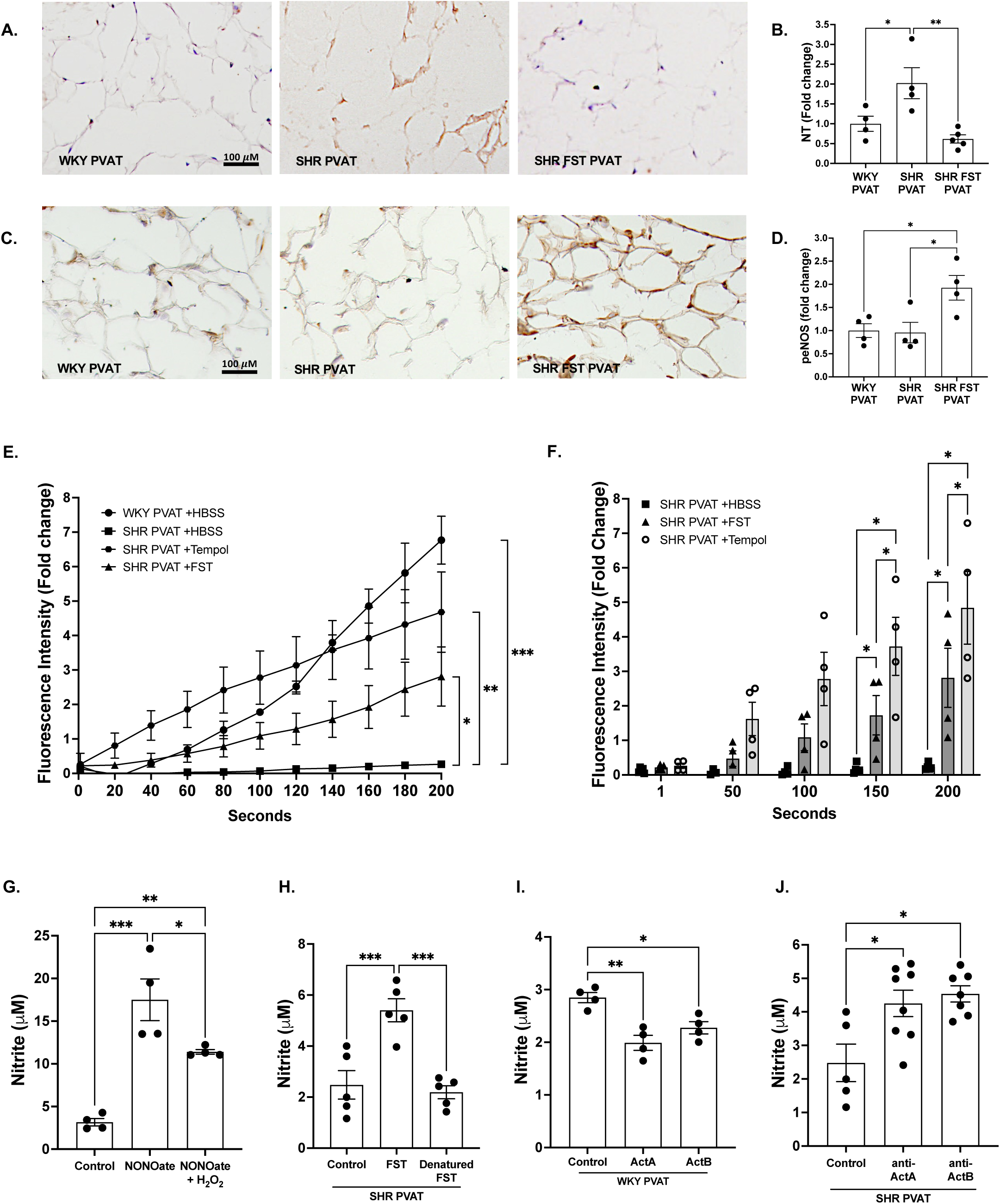
Follistatin increases NO bioavailability in SHR PVAT. Representative cross-sectional images of mesenteric PVAT from chronically treated rats stained for **A.** nitrotyrosine and **C.** phosphorylated eNOS, quantified in the accompanying graphs **B.**, and **D. E.** Fluorescence intensity measured over a 200-second period in mesenteric PVAT treated for 30 minutes with HBSS, follistatin or Tempol stained with DAF2-FM. **F.** Data in **E.** reported as individual values taken every 50 seconds. **G.** Nitrite levels measured in PBS solution, NONOate and/or H_2_O_2_. **H.** Nitrite levels measured in lysate of SHR PVAT treated for 30 minutes with follistatin or heat-denatured follistatin. **I.** Nitrite levels measured in lysate of WKY PVAT treated for 30 minutes with activin A or B. **J.** Nitrite levels measured in lysate of SHR PVAT treated for 30 minutes with neutralizing antibodies to activin A or B. *\*p*<0.05, *\*\*p* <0.01, *\*\*\*p* <0.001.

We next assessed whether H_2_O_2_ reduced NO in PVAT. Under physiological conditions, NO is rapidly oxidized to nitrite, measured fluorometrically by the Greiss reagent. In PBS alone, NONOate-generated nitrite was reduced by addition of H_2_O_2_ **(Figure 3G)**. **Figure 3H** shows that follistatin, but not heat denatured follistatin, increases nitrite production in SHR PVAT. These data suggest that the biological function of follistatin is required for its increase of NO availability. In keeping with this, activins A and B reduce nitrite levels in WKY PVAT **(Figure 3I)**. Conversely, activin A or B neutralization increase nitrite production in SHR PVAT **(Figure 3J)** highlighting a role for activins in regulating NO likely via ROS.

### Follistatin induces adipose tissue browning

Adipose tissue browning is associated with improved vasorelaxation and more recently, lower BP^9^. In both *in vitro* and *in vivo* models, follistatin was shown to induce peripheral adipose depot browning, although PVAT was not studied. We thus assessed whether follistatin induces browning of white PVAT. We first determined adipocyte size which is smaller in brown compared with white adipose tissue^23^. **Figure 4A, B** shows that PVAT adipocytes were larger in SHR compared with WKY, with size reduced in SHR by follistatin. This was accompanied by increased expression of the BAT markers PRDM16 **(Figure 4C, D)** and UCP-1 **(Figure 4E, F)**. In periaortic PVAT which has a brown phenotype, the downregulation of UCP-1 in SHR was reversed by follistatin **(Figure 4G, H)**. These data show that follistatin can induce browning in both phenotypically white and brown PVAT. We confirmed that follistatin also induced browning of a peripheral WAT depot, inguinal WAT, evidenced by increased PRDM16 **(Figure 4I, J)**.

**Figure 4:**
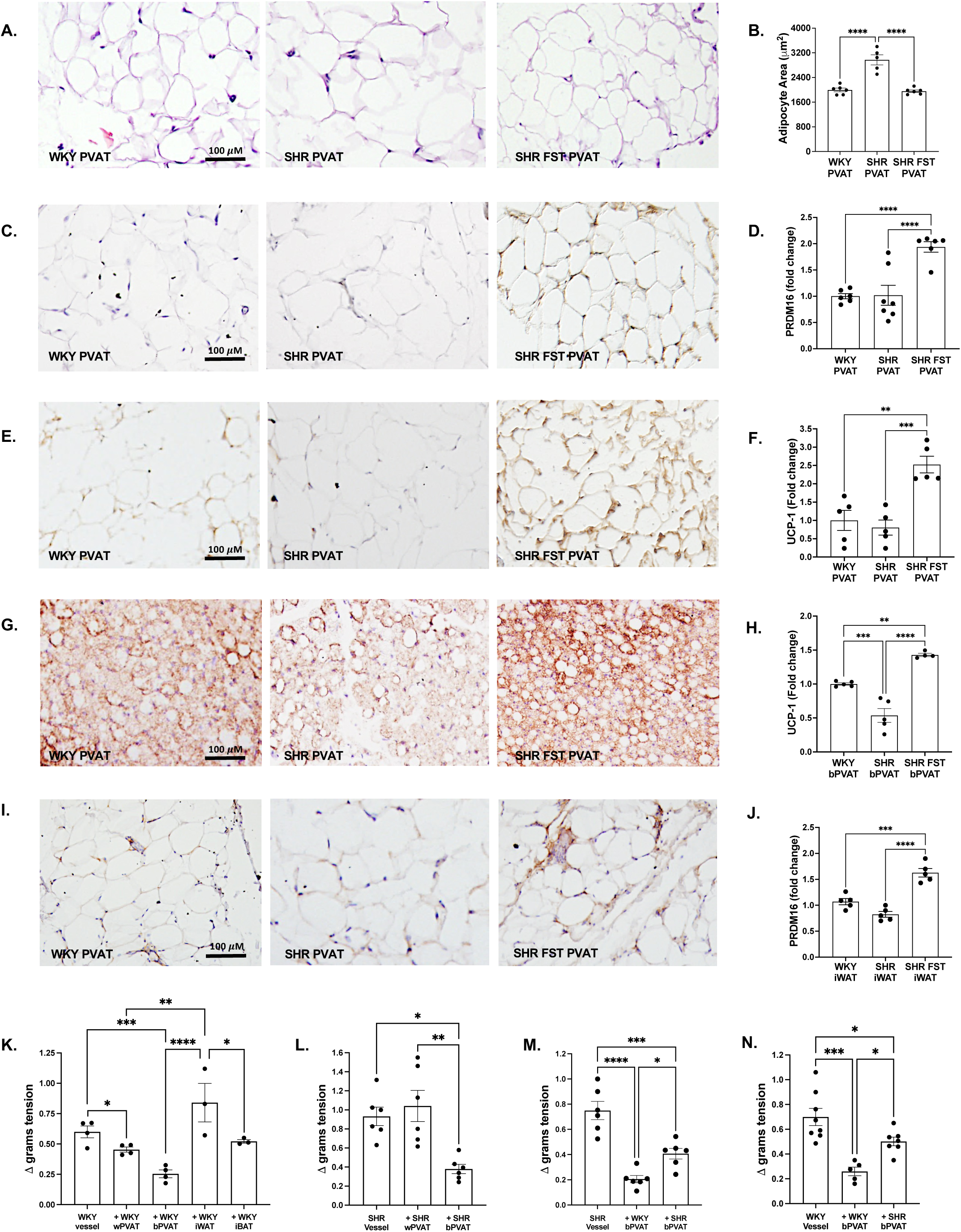
Follistatin induces characteristics of brown adipose tissue associated with vasorelaxant properties. Representative cross-sectional images of mesenteric PVAT from chronically treated rats stained for **A.** hematoxylin and eosin, **C.** PRDM16, **E.** UCP-1, quantified in accompanying graphs **B., D.**, and **F.** Representative cross-sectional images of **G.** thoracic aorta PVAT and **I.** inguinal WAT from chronically treated rats stained for UCP-1 and PRDM16 respectively and quantified in accompanying graphs **H.**, and **J. K.** Response to maximal KCl dose (125 mM) in first-order WKY mesenteric vessels preincubated with either detached WKY white mesenteric PVAT (wPVAT), brown thoracic aorta PVAT (bPVAT), inguinal WAT (iWAT) or interscapular BAT (iBAT). **L.** Response to maximal KCl dose (125 mM) in first-order SHR mesenteric vessels preincubated with either detached SHR wPVAT or bPVAT. **M.** Response to maximal KCl dose (125 mM) in first-order SHR mesenteric vessels preincubated with either detached WKY bPVAT or SHR bPVAT. **N.** Response to maximal KCl dose (125 mM) in first-order WKY mesenteric vessels preincubated with either detached WKY bPVAT or SHR bPVAT. Contractility values are from 3-5 rats per group, 1-3 vessels from each rat. *\*p*<0.05, *\*\*p* <0.01, *\*\*\*p* <0.001, *\*\*\*\*p* <0.0001.

We next determined whether white and brown PVAT have different effects on vessel response to KCl. WKY vessels were incubated with detached white mesenteric or brown aortic PVAT and constriction to KCl was assessed. As shown in **Figure 4K**, brown PVAT induced a more potent anticontractile effect in WKY vessels compared to white PVAT. Interestingly, WAT from a peripheral depot (inguinal) did not modulate contraction, highlighting an important functional difference between peripheral depots and PVAT.

We next determined whether brown PVAT from SHRs showed impaired anticontractile effects similar to white PVAT. We performed detached studies in which brown or white SHR PVAT was detached and incubated with an SHR vessel without intact PVAT, and constriction was induced using KCl. As seen in **Figure 4L**, detached SHR brown but not white PVAT resulted in an anticontractile effect on SHR vessels, suggesting that a phenotypically brown PVAT may improve vascular contraction. SHR vessels demonstrated a more pronounced decrease in contraction upon coincubation with WKY brown PVAT, which was significantly less than the constriction induced by SHR brown PVAT **(Figure 4M)**. These findings suggest improved regulation of vascular function with PVAT from non-hypertensive animals. To determine whether the blunted anticontractile effect of SHR brown PVAT was a result of impaired vessel or PVAT function, detached brown PVAT was incubated with WKY vessels. The anticontractile effect of SHR brown PVAT was reduced compared with WKY brown PVAT, highlighting dysfunction in SHR PVAT itself **(Figure 4N)**. Together these results illustrate potent vasodilatory effects of phenotypically brown PVAT which is reduced in the SHR.

### AMPK regulates follistatin-mediated PVAT browning

*In vivo*, follistatin has been shown to induce browning of peripheral adipose tissue depots via increased phosphorylation and activation of ERK 1/2^16^, pp38 MAPK^17^ and AMPK^18^. Here we demonstrate that follistatin did not affect activation of ERK 1/2 **(Figure 5A, B)**, but increased phosphorylation of AMPK **(Figure 5C, D)** in SHR PVAT. Follistatin also increased phosphorylation of pp38 MAPK **(Supplementary Figure 2A, B)**.

**Figure 5:**
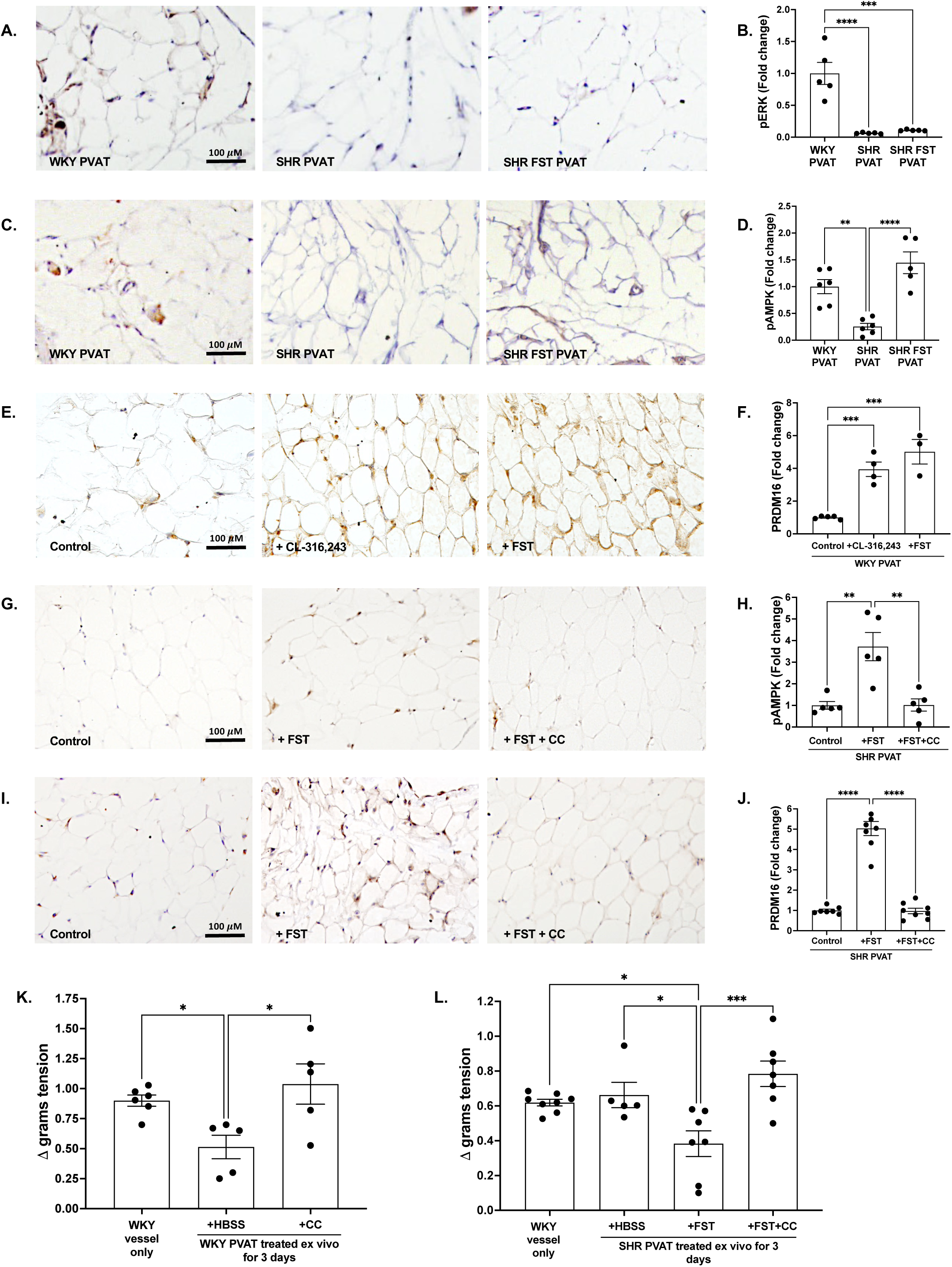
Follistatin induces PVAT browning via AMPK. Representative cross-sectional images of mesenteric PVAT from chronically treated rats stained for **A.** phosphorylated ERK and **C.** phosphorylated AMPK, quantified in accompanying graphs **B.** and **D. E.** Representative cross-sectional images of WKY mesenteric PVAT treated ex vivo for 3 days with either CL-316,243 or follistatin and stained for PRDM16, quantified in **F. G.** Representative cross-sectional images of SHR mesenteric PVAT treated ex vivo for 3 days with follistatin with or without Compound C (CC) and stained for phosphorylated AMPK, quantified in **H. I.** Representative cross-sectional images of SHR mesenteric PVAT treated ex vivo for 3 days with follistatin with or without CC and stained for PRDM16, quantified in **J. K.** Response to maximal KCl dose (125 mM) in WKY vessels incubated with mesenteric WKY PVAT treated ex vivo for 3 days with either HBSS or CC. **L.** Response to maximal KCl dose (125 mM) in WKY vessels incubated with mesenteric SHR PVAT treated ex vivo for 3 days with either HBSS, follistatin or follistatin and CC. Contractility values are from 2-3 rats per group, 2-3 vessels each rat. *\*p*<0.05, *\*\*p* <0.01, *\*\*\*p* <0.001, *\*\*\*\*p* <0.0001.

To further study the mechanism by which follistatin induces PVAT browning, we established an ex vivo culture system for PVAT as described by Blumenfeld et al^24^. Fresh mesenteric white PVAT was isolated and cultured in 30 mg sections for 3 days. We first confirmed that browning could be established in this system by treating isolated PVAT with the β_3_ adrenergic receptor agonist CL-316,243, a known inducer of browning in vivo^25^. CL-316,243 and follistatin increased PRDM16 in WKY PVAT **(Figure 5E, F)**. Ex vivo 3-day follistatin treatment in SHR PVAT increased phosphorylated AMPK, which was inhibited by cotreatment with the AMPK inhibitor compound C **(Figure 5G, H)**. Follistatin also induced browning evidenced by increased PRDM16 expression **(Figure 5I, J)**. This was inhibited by compound C, suggesting that follistatin-induced PVAT browning in the SHR is dependent on AMPK.

We next determined the role of AMPK in PVAT regulation of vascular function. As shown in **Figure 5K**, treatment of WKY PVAT with compound C for 3 days prior to incubation with vessels for myography abolished its anticontractile effect to KCl. Conversely, treating SHR PVAT ex vivo for 3 days with follistatin improved its ability to induce vasodilation in WKY vessels, an effect which was eliminated by cotreatment of SHR PVAT with compound C **(Figure 5L)**. Taken together, these results demonstrate that follistatin induces PVAT browning in SHRs through AMPK activation which contributes to its anticontractile effect on vessels.

### Activin A or B neutralization induces PVAT browning

As follistatin is a potent inhibitor of activins A and B, we next assessed the involvement of each of these to PVAT browning using neutralizing antibodies. Activin A or B neutralization each induced browning in SHR PVAT as evidenced by increased PRDM16 expression **(Figure 6A, B)**. On the contrary, treating WKY PVAT with recombinant activin A or B reduced PRDM16 expression **(Figure 6C, D)**. Knockout of Smad3, a major downstream activin signalling mediator, was shown to increase browning of peripheral WAT ^26^. As increased Smad3 phosphorylation was also shown to inhibit AMPK activity^27^ we sought to determine whether Smad3 inhibition would increase AMPK phosphorylation/activation. The Smad3 inhibitor SIS3, increased AMPK activation in the presence of either activin A or B (**Supplementary Figure 3A-D**). These results suggest that activin signalling regulates PVAT browning via Smad3 regulation of AMPK.

**Figure 6:**
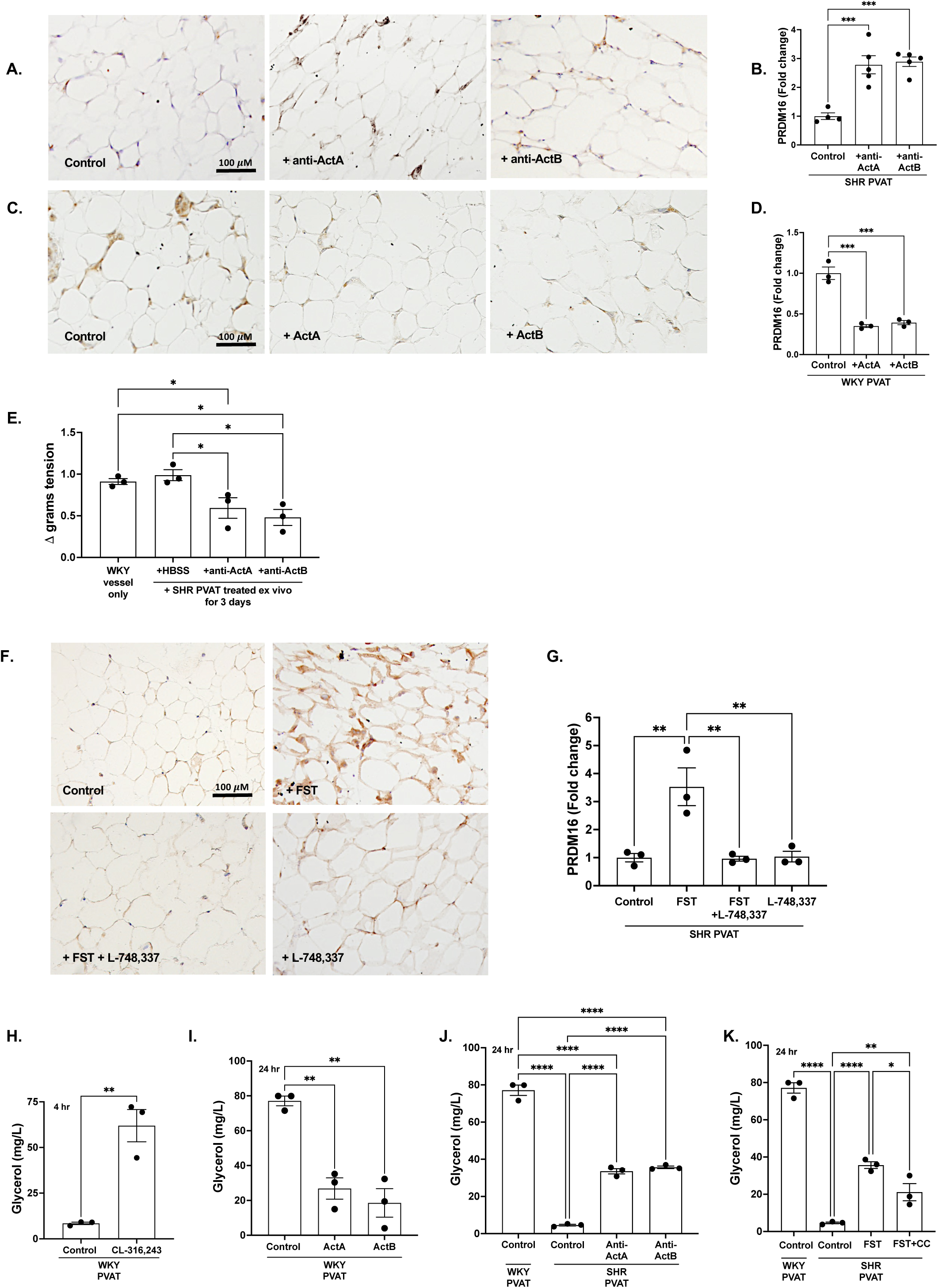
Follistatin induces PVAT browning and lipolysis via neutralization of activins and β_3_ receptor activation. **A.** Representative cross-sectional images of SHR mesenteric PVAT treated ex vivo for 3 days with either HBSS or neutralizing antibodies to activin A or activin B and stained for PRDM16, quantified in **B. C.** Representative cross-sectional images of WKY mesenteric PVAT treated ex vivo for 3 days with either HBSS, activin A or activin B and stained for PRDM16, quantified in **D. E.** Response to maximal KCl dose (125 mM) in WKY vessels incubated with mesenteric SHR PVAT treated ex vivo for 3 days with either HBSS or neutralizing antibodies to activin A or activin B. **F.** Representative cross-sectional images of SHR mesenteric PVAT treated ex vivo for 3 days with either HBSS, follistatin, follistatin and L-748,337 or L-748,337 alone, quantified in **G. H.** Glycerol content measured in media from WKY PVAT treated with either HBSS or CL-316,243 for 4 hours. I. Glycerol content measured in media from WKY PVAT treated with either HBSS, activin A or activin B for 24 hours. **J.** Glycerol content measured in media from SHR PVAT treated with either HBSS or neutralizing antibodies to activin A or activin B for 24 hours. **K.** Glycerol content measured in media from SHR PVAT treated with either HBSS, follistatin or follistatin and CC for 24 hours. *\*p*<0.05, *\*\*p* <0.01, *\*\*\*p* <0.001, *\*\*\*\*p* <0.0001.

We next sought to determine whether activin inhibition in SHR PVAT could restore its vasodilatory effects. SHR PVAT treated with activin A or B neutralizing antibodies for 3 days ex vivo improved PVAT ability to reduce vasoconstriction to KCl in WKY vessels **(Figure 6E)**. Conversely, treating WKY PVAT acutely (30 minutes) with activin A or B reduced its anticontractile effect **(Supplementary Figure 4)**. These results suggest that activins decrease adipose tissue browning to contribute to PVAT dysfunction in SHRs.

Lastly, we assessed the interaction between ROS, activin inhibition and AMPK activation in promoting PVAT browning. Inducing browning in SHR PVAT with CL-316,243 significantly reduced ROS **(Supplementary Figure 5A, B)**. Conversely, ROS inhibition in SHR PVAT increased PVAT browning which was attenuated by AMPK inhibition **(Supplementary Figure 5C, D)**, suggesting that ROS regulates browning via AMPK. Indeed, AMPK inhibition also increased ROS in WKY PVAT **(Supplementary Figure 5E, F)**. Together, these results suggest that follistatin induces SHR PVAT browning via activin neutralization, ROS inhibition and AMPK activation.

### Follistatin enhances response to β_3_ receptor activation

A well-known mechanism of adipose tissue browning involves activation of β_3_ adrenergic receptors^28^. To determine whether β_3_ adrenergic receptors are required for follistatin-induced browning, we used the β_3_ blocker L-748,337. Follistatin-induced browning of SHR PVAT as assessed by PRDM16 induction was reduced by β_3_ receptor inhibition **(Figure 6F, G)**. β_3_ adrenergic receptor inhibition alone had no effect. Together these results suggest that PVAT browning induced by follistatin requires activation of β_3_ adrenergic receptors.

β_3_ adrenergic receptor activation is known to stimulate lipolysis, producing free fatty acids that can upregulate and activate UCP-1^28^. Thus, we next determined whether follistatin induced lipolysis in SHR PVAT. The release of glycerol, a byproduct of lipolysis, was measured in media collected from cultured PVAT. We first confirmed that CL-316,243 could increase glycerol secretion, seen after 4 hours **(Figure 6H)**. Activin A or B significantly reduced glycerol in cultured media from WKY PVAT, indicative of supressed lipolysis **(Figure 6I)**. Conversely, activin A or B neutralization or follistatin significantly increased lipolysis in SHR PVAT **(Figure 6J, K)**. Follistatin effects were reduced by AMPK inhibition **(Figure 6K)**. These results demonstrate that follistatin induces lipolysis in SHR PVAT via activin inhibition and AMPK activation, suggesting an additional potential mechanism of increased browning.

### BP lowering alone does not improve vascular function or induce adipose tissue browning

Studies have correlated adipose tissue browning with improved vascular function and lower BP^9^. Whether reducing elevated BP results in adipose tissue browning has not been explored. Here, we assessed the effects of the vasodilator hydralazine on vascular function, targeting a similar BP reduction which we achieved previously with follistatin. Rats were treated for 8 weeks. As shown in **Supplementary Figure 6A, B**, hydralazine lowered systolic and diastolic BP to a similar extent as follistatin. Unlike that seen with follistatin, however, hydralazine did not reverse the procontractile effect of SHR PVAT **(Supplementary Figure 6C)**, nor did it induce adipose tissue browning as measured by PRDM16 (**Supplementary Figure 6D, E)**. Altogether, BP lowering alone neither improves vascular function nor induces adipose tissue browning in the SHR, suggesting that follistatin directly affects PVAT and vascular function to improve BP.

### Follistatin shifts the SHR PVAT proteome towards WKY

Unbiased proteomic analysis of mesenteric PVAT from rats treated for 8 weeks with follistatin or vehicle was performed using LC-MS to investigate differential protein expression and processes regulated by follistatin compared to SHR controls **(Figure 7A)**. A total of 3208 proteins were identified in the sample with 2822 of these having high false discovery rate (FDR) confidence. Proteins with low and medium FDR confidence were not included in the analysis. An additional 64 contaminant proteins (2.88% of total) were identified in the samples and were excluded from the analysis. Using the SHR vehicle as denominator, 24 proteins were down-regulated in the SHR follistatin group and 30 proteins were up-regulated **(Supplementary Figure 7)**. Principal component analysis (PCA) plots revealed a distinguishable difference between vehicle-and follistatin-treated SHRs **(Figure 7B)**, with direction of the follistatin-treated SHR cluster resembling that of WKY controls. Heatmap analysis of protein abundances highlights distinct profiles between vehicle and follistatin-treated SHRs **(Figure 7C)**.

**Figure 7:**
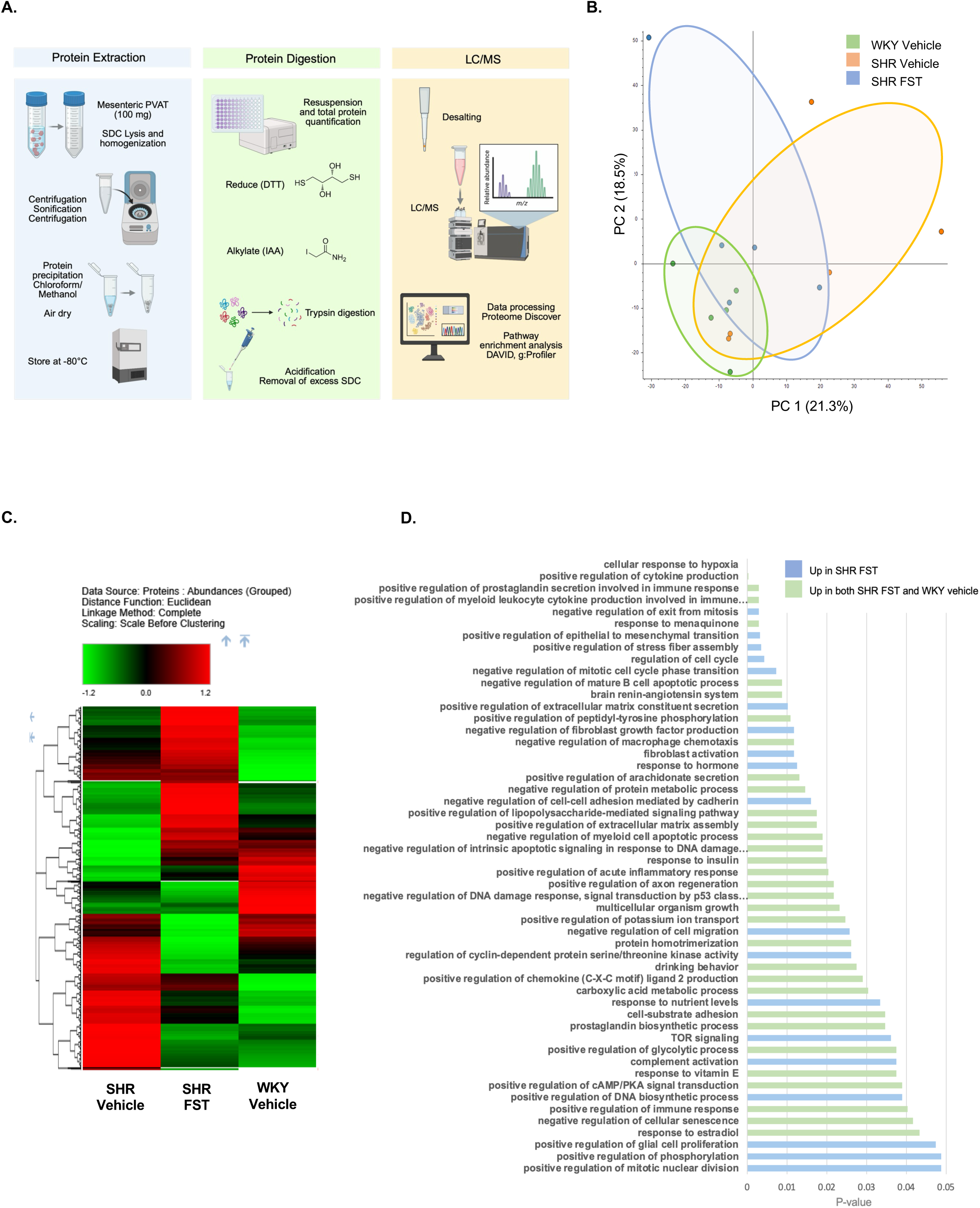
Analysis of differentially expressed proteins in mesenteric PVAT by mass spectrometry. **A.** Schematic of PVAT sample preparation and processing for proteomic analysis. **B.** Principal component analysis of PVAT proteome from the 3 chronic treatment groups: WKY vehicle (green), SHR vehicle (yellow) and SHR follistatin (blue). **C.** Heatmap of differentially expressed proteins. **D.** Gene ontology of differentially expressed proteins by biological processes, upregulated in PVAT from both WKY control and follistatin-treated SHRs (green) and follistatin-treated SHRs alone (blue) in comparison to SHR controls. *p* values are indicated in the x-axis of bar graphs. SDC indicates sodium deoxycholate; DTT, dithiothreitol; IAA, iodoacetamide; LC/MS, liquid chromatography/mass spectrometry.

Protein abundances from follistatin-treated SHRs align more closely to WKY controls. Gene Ontology analysis revealed 52 biological processes significantly upregulated in PVAT by follistatin compared to SHR PVAT **(Figure 7D)**, with 63% of these also upregulated in WKY PVAT compared to SHR PVAT. Follistatin increased processes that regulate adipose tissue browning including cAMP, PKA signalling and that may induce vasodilation. Collectively, these data reveal that follistatin induces a shift in the SHR PVAT proteome toward a normal PVAT profile.

## Discussion

Impairment in PVAT, which normally functions to exert an overall anticontractile effect, is increasingly recognized as an important contributor to hypertension. In this study, we identify impaired regulation of vascular tone by SHR PVAT. Follistatin restores PVAT function by inhibiting oxidative stress and inducing browning, mediated by neutralization of activins A and B and AMPK activation **(Figure 8)**. This is the first study to assess the effects of PVAT oxidative stress and browning on resistance artery function in hypertension, identifying follistatin as a novel regulator restoring PVAT function.

**Figure 8:**
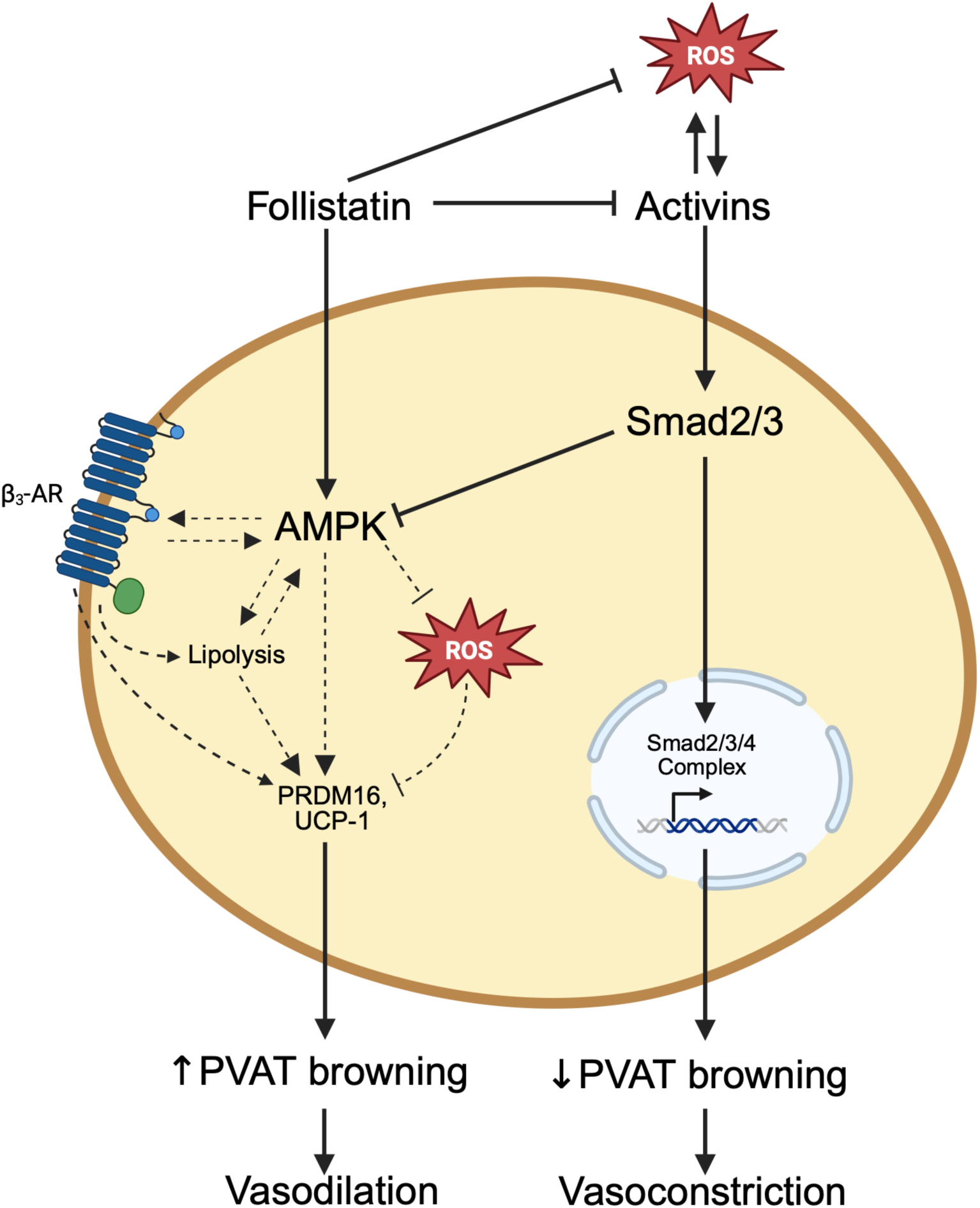
Schematic summary. Follistatin restores the anticontractile effect of SHR PVAT through the inhibition of oxidative stress and induction of adipose tissue browning. This is mediated by activin neutralization and AMPK phosphorylation. Improved PVAT function is associated with reduced BP in the SHR model of essential hypertension. Schematic diagram designed using BioRender.

We had previously identified that SHR vessels exhibit increased oxidative stress, and this is reduced by follistatin in association with improved vessel function^10^. Although the mesenteric PVAT anticontractile effect is known to be impaired in this model^29^, the effect of follistatin on PVAT was unknown. We now show that the increased oxidative stress seen in SHR PVAT, known to contribute to its dysfunction^30^, is abolished by both chronic in vivo and acute ex vivo follistatin treatment. This reduction in oxidative stress by follistatin has also been observed in other settings such as chronic kidney disease from reduced kidney mass^31^. Interestingly, our data further highlight the ability of PVAT to induce oxidative stress to a closely associated vessel, an effect which is prevented by follistatin. Indeed, the ability of PVAT-generated ROS to diffuse towards vascular cells and cause oxidative damage and impaired vessel function is known^32^. Follistatin thus has dual protective effects on the vessel in hypertensive animals which is both direct and indirect through reducing PVAT oxidative stress.

Follistatin neutralizes several members of the TGFβ superfamily, with greatest potency against activins A and B. We and others have reported that activin A contributes to vessel ROS production in both systemic and pregnancy-associated hypertension (preeclampsia)^10, 12^. However, the contribution of activin B to oxidative stress in general, as well as the contribution of activins A and B to PVAT oxidative stress and dysfunction in hypertension were unknown. We demonstrate increased abundance of activins A and B in SHR PVAT, both of which were reduced by follistatin. Activin A or B treatment of normal PVAT augmented ROS production, suggesting that activin signalling directly contributes to ROS production in PVAT. While ROS production in response to activin A was mediated by its induction of NADPH oxidase expression in vessels^33^, whether this mechanism is relevant to PVAT remains to be explored.

ROS contribute to HTN through reduction in bioavailability of the vasodilator, NO^34^. PVAT is a major producer of NO which is known to be reduced in SHR PVAT^30^. Here we show that follistatin improves NO bioavailability in SHR PVAT, mediated by both activins A and B. Within PVAT, NO is produced primarily by endothelial NO synthase (eNOS). Activation of eNOS, as measured by its phosphorylation at S1177, was not different between WKY and SHR PVAT, in keeping with findings by Carvalho et al ^30^. However, follistatin increased eNOS activation in SHR PVAT, suggesting that the restored NO bioavailability may be due both to reduced scavenging by superoxide as well as increased production.

In individuals, the presence of BAT correlates with lower prevalence of hypertension^8^, suggesting that increased browning of PVAT may promote its anticontractile effects. Indeed, in models of obesity and HTN, reduced PVAT browning or increased whitening reduced its anti-contractile properties. Conversely, β_3_ agonist-induced browning of PVAT increased its anticontractile effect^34^. Here we show that follistatin induced browning of peripheral adipose depots, as seen previously ^16,17,18^, and now for the first time, demonstrate that follistatin also induces browning of PVAT. Furthermore, we show that brown PVAT provides a more potent anticontractile effect compared to white PVAT. We additionally highlight a functional difference between peripheral adipose tissue depots and PVAT, such that PVAT is more anticontractile than peripheral depots, even within the same phenotype. Heterogeneity in the secretion profiles of vasoactive factors and activity of the sympathetic nervous system are likely contributors to these differences.

While follistatin did not alter ERK activation in our study, both p38 MAPK and AMPK activation were reduced in SHR PVAT and increased by follistatin. As AMPK deficiency results in suppressed PVAT NO synthesis and impaired anticontractile activity^35^, we elected to focus further on its assessment. We identified that PVAT browning and its anticontractile properties are dependent on AMPK activity. While our studies did not address the mechanism by which follistatin increases AMPK activation, elevated AMP levels as seen in response to follistatin in mouse embryonic fibroblasts may have contributed^16^.

Our studies identified that activin neutralization increased PVAT browning ex vivo. Whether activin signaling reduces AMPK activity is unknown, but additional mechanisms by which activins could reduce browning have been identified. In cultured adipocytes, activin A reduced expression of BAT genes^36^. Conversely, blocking activin signaling with a type 2 receptor inhibitor suppressed Smad3 signalling to increase brown adipocyte differentiation^37^. Interestingly, AMPK was shown to attenuate Smad2/3 activation^27^, suggesting that it may inhibit activin signaling to promote PVAT browning.

β_3_ adrenergic receptor activation is an important stimulator of browning through its induction of lipolysis and consequent release of free fatty acids that bind to and activate UCP-1. Activin B and Smad2/3 signaling were shown to inhibit lipolysis in adipocytes^38^, suggesting that activins may inhibit browning. In agreement, we found reduced lipolytic activity in SHR PVAT which was improved with follistatin and both activin A and B neutralization. The improved lipolysis with follistatin was reduced by inhibition of AMPK, a well-known mediator of β_3_ adrenergic effects^39^. Interestingly, Singh et al demonstrated that peripheral adipose depot browning in response to CL-316,243 was enhanced in follistatin transgenic mice^17^. While AMPK was not investigated, this effect was due at least in part to p38 activation which we showed to also be restored by follistatin. As p38 is known to be activated by AMPK in adipocytes, further studies would address whether follistatin restores p38 activity via activation of AMPK.

Intersection between browning, oxidative stress and AMPK has been suggested. Browning was shown to reduce oxidative stress^40^, in keeping with our ex vivo studies in which induction of PVAT browning reduced oxidative stress in SHR PVAT. Conversely, reduction in BAT through UCP-1 deficiency resulted in increased oxidative stress and elevated BP^41^, associated with reduced AMPK activation^42^. Our findings that AMPK inhibition increases PVAT ROS in association with a reduced brown phenotype highlight the complex interplay between browning, ROS and AMPK activity.

Proteomic analysis of mesenteric PVAT broadly showed that follistatin shifted the proteome toward characteristics of normal PVAT. Pathway enrichment analysis identified processes associated with cytokine secretion, cell cycle regulation, extracellular matrix, immune response and prostaglandin synthesis as being restored by follistatin, in keeping with the multicellular composition and sophisticated environment that constitutes PVAT. While these data support mechanisms regulating browning, including cAMP and PKA signalling, future studies would assess additional processes in more detail. The following are worth noting: 1) Follistatin increased synthesis of prostaglandins which may induce vasodilation, such as those signaling through E-prostanoid receptors 2 and 4 (EP_2_, EP_4_)^43^. Indeed, EP_2_-induced vasodilation is known to be reduced in SHR^44^. Furthermore, the enzyme cyclooxygenase, a proximal regulator of prostaglandin synthesis, is required for cold-induced browning of inguinal WAT^45^. 2) Increased apoptosis also characterizes PVAT dysfunction in the SHR^30^. One of the most upregulated proteins by follistatin was the anti-apoptotic protein CIAPIN1 (cytokine-induced apoptosis inhibitor 1), suggesting that follistatin may also protect against this process.

This study has several limitations that should be acknowledged. First, only male animals were included in the analysis, and therefore potential sex-specific differences cannot be assessed. Given known sex differences in hypertension, future studies incorporating female animals will be important to determine generalizability of these findings. Second, PVAT is heterogeneous, producing a wide range of bioactive mediators. Proteomic analysis would not capture non-protein factors such as gaseous mediators (for example, hydrogen sulfide) or lipid-derived signaling molecules, which may contribute to vascular regulation. In addition, our proteomic analysis is inherently limited by the availability and annotation of characterized proteins within the rat proteome database, which may restrict the identification of relevant targets.

In summary, our findings establish follistatin as a potent modulator of PVAT function, capable of preventing its procontractile effect through activin neutralization and AMPK-dependent browning, and attenuation of oxidative stress. The improved vascular function is associated with lower BP in the SHR model of essential hypertension. These insights underscore the role of PVAT in vascular homeostasis, advancing our understanding of adipose-vascular crosstalk. Our findings provide a mechanistic foundation for the development of PVAT-targeted therapies to mitigate vascular dysfunction in essential hypertension.

## Acknowledgements

The authors acknowledge support from The Research Institute of St. Joe’s for nephrology research. They are also grateful to Paranta Biosciences Limited (Australia) for providing human recombinant FST-288.

## Supplementary Material

Supplementary Methods

Figure S1-S7

References 10, 19, 20

## Supplementary Methods

### Animal studies

Experiments adhered to guidelines set by McMaster University and the Canadian Council on Animal Care (CCAC) and were approved by the McMaster University Animal Research Ethics Board (#23-35). Research also adheres to ARRIVE guidelines. Male SHRs were utilized as a model of essential hypertension, while Wistar Kyoto (WKY) rats served as normotensive controls. Both rat strains were sourced from our in-house breeding colony. All tissues from chronic vehicle- or follistatin-treated rats used for experiments in this study are from our previously published work^10^. Briefly, male rats 12 weeks of age were divided into three groups: WKY Vehicle (n = 10), SHR Vehicle (n = 10), and SHR FST (follistatin, n = 10). Radiotelemeters (Data Sciences International) were implanted into the abdominal aorta as previously described^10^. Concisely, rats were anesthetized, and the BP-sensing catheter was inserted into the abdominal aorta below the renal arteries. The insertion site was sealed, and the transmitter was placed within the peritoneal cavity. After surgery, rats were housed individually and following one week of recovery, BP was transmitted, transduced via receivers (RPC-1; Data Sciences International) and recorded (Data Acquisition (DAQ)) for 2 hours weekly at noon.

Baseline BP of no less than 160/100 mmHg was required in SHRs for inclusion in the study. All SHRs met this requirement, and none were excluded. DAQ waveforms were routinely observed to confirm unobstructed blood flow of telemeter catheters. For hydralazine-treated SHRs, water bottles containing dissolved hydralazine were replaced every 3 days. Following baseline BP recording for 3 days, rats received intraperitoneal injections of either vehicle (PBS) or follistatin-288 (Paranta Biosciences, 0.075 mg/kg) every other day for eight weeks. In a second study for this manuscript, SHRs aged 12 weeks were treated with hydralazine (10 mg/kg/day) in drinking water for 8 weeks. The dose was selected based on its efficacy to reduce BP to the same extent as follistatin (20 mmHg BP reduction) from preliminary trials. All animals were weighed and appearance and body condition were assessed every other day prior to treatment administration.

### End-point sampling

At the end of the treatment period, rats were anesthetized using isoflurane and telemeters were carefully removed. An incision was made in the right atrium, and the left ventricle was punctured and perfused with 30 mL of Hank’s Buffered Saline Solution (HBSS, 0.14 M NaCl, 5.4 mM KCl, 0.25 mM Na_2_HPO_4_, 0.1 g glucose, 0.44 mM KH_2_PO_4_, 1.3 mM CaCl_2_, 1.0 mM MgSO_4_, 4.2 mM NaHCO_3_). A ligature was placed at the midpoint of the mesenteric bed. The distal half of the mesenteric bed was excised to isolate PVAT for liquid nitrogen storage as well as the vessels with intact PVAT for myograph contractility studies. Isolation was done under a dissecting microscope. Subsequently, the left ventricle was perfused with sodium nitroprusside (10^-6^ M) to achieve maximal dilation of the proximal half of the mesenteric bed. The animals were then perfused with 4% paraformaldehyde, after which the proximal half was removed. The dilated, fixed vessels were isolated and processed for histological analysis. First-order mesenteric arteries, thoracic aorta and the surrounding PVAT were utilized.

### Myograph contractility studies

Vessel contractility was assessed in first-order mesenteric arteries with or without PVAT using a wire myograph as previously described^10^. Briefly, vessels were cut into 2 mm rings prior to mounting with baseline tension set to 0.5 g and continuous aeration with 95% O_2_ and 5% CO_2_. Vessels were challenged with increasing doses of KCl (30 - 130 mM) or the β_1_ adrenergic receptor agonist phenylephrine (10^-8^ - 3×10^-5^ M) to elicit smooth muscle depolarization or receptor-mediated contraction respectively.

Contractility data were recorded using WINDAQ Data Acquisition software through a DI-720-USB Series analog-to-digital converter (DATAQ Instruments Inc). For detached PVAT studies, either white PVAT (surrounding first-order mesenteric arteries) or brown PVAT (surrounding thoracic aorta) were detached from the vessels and positioned next to mounted mesenteric vessels in the myograph chamber. Mounted vessels and detached PVAT were incubated together for 30 minutes after which the PVAT was removed, and the vessels were challenged with a maximal dose of KCl (125 mM). In strain crossover studies, vessels or PVAT were paired with tissues from the opposite strain. All acute ex vivo treatments of detached PVAT were conducted in 24-well plates at 37℃ for 30 minutes prior to co-incubation with vessels in the myograph chamber for an additional 30 minutes. To eliminate the contribution of the endothelium, this layer was denuded using human hair. Complete endothelium removal was confirmed in vessels pre-constricted with phenylephrine (10^-6^ M) by assessing relaxation to carbachol, an endothelium-dependent acetylcholine-mimetic.

### Oxidative stress assessment

ROS were visualized in PVAT using the oxidation-sensitive fluorescent dye dihydroethidium (DHE, Molecular Probes) as previously described^10^. Wells from a 96 well-plate were coated with 1 μL of tissue adhesive (Corning Cell Tak) to adhere PVAT to the bottom. Fresh PVAT (<1 cm) taken from rats enrolled in treatment studies was placed at the centre of each well and incubated in HBSS and DHE (50 #M) at 37°C for 30 minutes. For ex vivo treatment, fresh PVAT from untreated rats was isolated and treated with either follistatin-288 (500 #M, Paranta Biosciences), Tempol (10 #M, Sigma-Aldrich), activin A (50 ng/ml, R&D Systems), activin B (50 ng/ml, R&D Systems), SB-431,542 (5 #M, Caymen Chemicals) or DMSO for 30 minutes prior to incubation with DHE for an additional 30 minutes. In some cases, PVAT were treated ex vivo for 3 days followed by DHE at 37°C for 30 minutes prior to imagining. PVAT were imaged using a fluorescent microscope (Nikon Eclipse Ti-E, NIS-Elements AR program) with excitation 510 nm, emission 595 nm. Images of DHE-stained PVAT were captured using Metamorph software and analyzed using ImageJ.

### Immunohistochemistry

Paraformaldehyde-fixed PVAT was embedded in paraffin blocks and cut into 4 µm sections. Tissues were deparaffinized for immunohistochemical analysis. FST, activin A and B expression were assessed using antibodies to FST (1:200, Proteintech), Activin A (1:50, R&D Systems) and Activin B (1:1000, Abnova). The presence of peroxynitrite residues, a product of ROS and NO, was assessed using anti-nitrotyrosine (1:6000, Millipore). Adipocyte area was measured in PVAT tissues stained using hematoxylin and eosin (Sigma). PVAT browning was assessed in tissues using antibodies to PRDM16 (1:500, Millipore) and UCP-1 (1:500, Alpha Diagnostics). To describe specific mechanisms of follistatin-induced browning, PVAT were stained with pp38 MAPK (1:500, Cell Signalling), pErk 1/2 (1:500, Cell Signalling) and pAMPK Thr172 (1:200, Cell Signalling). To assess inflammatory markers in PVAT, tissues were stained with CD68 (1:200, Bio-Rad Antibodies), iNOS (1:500, Abcam) and CD206 (1:500). Staining was quantified using ImageJ.

### H_2_O_2_ measurements

H_2_O_2_ levels in WKY or SHR PVAT were assessed using the Amplex Red Hydrogen Peroxide/Peroxidase Assay Kit (Thermo Fisher Scientific). Approximately 60 mg of fresh mesenteric PVAT was dissected and weighed for each experimental condition. Tissues were incubated at 37℃ for 30 minutes in the presence of one of the following treatments: HBSS, pyocyanin (10 #M, Caymen Chemicals), follistatin (500 #M), heat-denatured follistatin, or activin A and B neutralizing antibodies (3.5 #g/mL). Following treatment, tissues were homogenized in HBSS using a bead homogenizer (Bead Ruptor Elite, Omni International) to ensure complete lysis. The resulting lysates were centrifuged twice at 15,000 x g for 15 minutes at 4℃. Supernatants were collected for H_2_O_2_ quantification following the manufacturer’s protocol. Briefly, each sample was incubated with Amplex Red reagent (10-acetyl-3,7-dihydroxyphenoxazine) and horseradish peroxidase in a 96-well plate. Fluorescence was measured at excitation 530 nm, emission 590 nm using a microplate reader after a 30-minute incubation at room temperature in the dark. H_2_O_2_ concentrations were calculated from a standard curve generated using known concentrations prepared in parallel.

### Direct NO detection

NO availability was determined using the cell-permeable fluorescent NO indicator, 4-Amino-5-Methylamino-2’,7’-Difluorofluorescein Diacetate (DAF-2 FM). After passive diffusion into the cell, it undergoes deacetylation by esterase which traps it intracellularly. Reaction with NO produces a highly fluorescent product. PVAT was isolated and adhered to the wells of a 96 well-plate, followed by incubation with DAF-2 FM (10 #M) for 1 hour. At the 30-minute mark, either follistatin (500 #M) or Tempol (10 #M) was added for 30 minutes at 37°C. Live imaging was performed using an Olympus IX81 fluorescence microscope equipped with a temperature-controlled stage maintained at 37℃. Baseline fluorescence was recorded for 20 seconds at 1 frame/second using excitation/emission of 495/515 nm. PVAT was then stimulated with the NO donor, NONOate (10 #M) and fluorescence accumulation was monitored for 200 seconds at the same acquisition rate. Image acquisition and quantitative analysis were conducted using Metamorph imaging software.

### Indirect NO detection

NO production was assessed indirectly by quantifying nitrite levels using the Griess Reagent Kit (Molecular Probes, Thermo Fisher Scientific). An initial assay was performed in PBS containing NONOate (10 #M) with or without H_2_O_2_, incubated for 30 minutes at 37℃. For tissue-based assays, 60 mg of mesenteric PVAT was isolated and incubated at 37℃ for 30 minutes with either follistatin (500 #M), heat-denatured follistatin, recombinant activin A or B (50 nM), or activin A or B neutralizing antibodies (3.5 #g/mL). Tissues were then homogenized in PBS using a bead homogenizer, and lysates were clarified by centrifugation at 15,000 x g for 10 minutes at 4℃. Supernatants were incubated with an equal volume of Griess reagent in a 96-well plate for 30 minutes at room temperature in the dark. Absorbance was measured at 540 nm using a microplate reader, and nitrite concentrations were calculated from a nitrite standard curve.

### PVAT browning ex vivo

This protocol was developed from a previous study^19^. Fresh mesenteric PVAT (60mg) was placed in 24-well plates containing Dulbecco’s Modified Eagle Medium (DMEM, Cat # 11965092) supplemented with 10% fetal bovine serum (FBS), 1% Penicillin/Streptomycin, 20 mM HEPES, 50 #g/mL sodium ascorbate and 1 #M insulin. PVAT was treated for 3 days with either CL-316,243 (2 #M, Cayman Chemicals), follistatin (500 #M), AMPK inhibitor Compound C (CC, 10 #M, Cayman Chemicals), activin A or B neutralizing antibodies (3.5 #g/mL), recombinant activin A or B (50 ng/mL), β_3_ adrenergic receptor blocker L-748,337 (100 #M, Cayman Chemicals), Tempol (10 #M) or selective Smad 3 inhibitor SIS3 (10 #M, Cayman Chemicals). On the third day, treated PVAT was fixed and stained by IHC.

### Lipolysis assay

Glycerol released by PVAT was used as a quantitative measure of lipolytic activity, based on a previous study^20^. White mesenteric PVAT was harvested, and 30 mg of tissue was weighed and cut into 5-6 segments. Tissue was placed in 48-well plates containing phenol red-free DMEM (Cat # 11965092) supplemented with 5% FBS at 37℃ for 15 minutes to establish baseline glycerol release. Subsequently, tissues were treated with either CL-316,243 (2 #M), recombinant activin A or B (50 ng/mL), activin A or B neutralizing antibodies (3.5 #g/mL), follistatin (500 #M) or Compound C (10 #M).

For CL-316,243-treated samples, conditioned media were collected at 4 hours post-treatment at 37℃. For all other treatment groups, media were collected after 24 hours at 37℃. Media were stored at 4℃ after collection. Glycerol concentrations in the collected media were quantified using a Cayman Chemical Lipolysis Assay Kit, following the manufacturer’s protocol.

### Sample preparation for adipose tissue proteomic analysis

Mesenteric PVAT was weighed to 100 mg and lysed in 4% sodium deoxycholate (SDC) prepared in Tris (1 M, pH 8.5). Tissue homogenization was performed using beads. Lysed samples were centrifuged twice (6000 x g at 4℃ for 15 minutes) and heated at 95℃ for 10 min. Samples were sonicated on ice using a probe sonicator (75% amplitude, 15:15s for 5 minutes). Samples were centrifuged and lysates were split into aliquots. Proteins were extracted using chloroform/methanol (2:1) in nuclease-free sterile water. Following top phase aspiration, replicates were centrifuged, combined and centrifuged again (7000 x g at 4℃ for 5 minutes). Remaining liquid was aspirated, and pellets were air dried and stored at -80℃ until ready for resuspension. Once 5 PVAT samples were processed for each treatment group, samples were randomized prior to transport to University Health Network for resuspension.

Pellets were resuspended in 1% SDC in ammonium bicarbonate (ABC). Lysates were sonicated in a water bath (5 x 5s on ice). Samples were centrifuged thrice (17000 x g at 4℃ for 10 minutes). Solubilized protein pellets from each aliquot per sample were pooled and the total protein content was measured using bicinchoninic acid (BCA) Protein Assay Kit (Thermo Fisher Scientific). Disulphide bonds were reduced with dithiothreitol (DTT, 10 mM) for 30 min at 56℃. Cysteine residues were alkylated with iodoacetamide (IAA, 20 mM) for 30 min at room temperature in the dark. Samples were reduced with DTT to remove excess IAA. CaCl_2_ (1mM) was added to each sample to stabilize trypsin. Each sample was digested with trypsin (20:1 protein to enzyme) overnight at 37℃. The following day, trifluoroacetic acid (TCA, 1%) was added to each sample to quench digestion and precipitate SDC.

### Liquid chromatography-mass spectrometry

Samples were processed by LC-MS by the McMaster Centre for Microbial Chemical Biology. Prior to desalting, pH was confirmed to be less than 2.0 for all samples. The samples were desalted using Peptide Desalting Columns (Thermo Fisher) and dried overnight on a lyophilizer (Labconco). The sample concentrations were measured using absorbance at 280 nm and samples then reconstituted in 0.1% (v/v) formic acid to a final peptide concentration of 0.5 #g/#l and transferred to polypropylene vials for LC-MS/MS. All samples were injected in triplicate. The peptides were analyzed by LC-MS/MS using an Ultimate 3000 RSLCnano HPLC system (Dionex) coupled to a Q Exactive HF mass-spectrometer (ThermoFisher). A volume of 4 #L was injected into a 75 #m i.d. x 75 cm Thermo Scientific Easy-Spray column packed with 2 #m resin. The LC run consisted of a 152-minute linear gradient of 3% to 35% acetonitrile followed by a 15-minute gradient from 35 to 50% acetonitrile and 10-minute gradient from 50 to 90% acetonitrile in 0.1% formic acid using a flow rate of 300 nL/min. The Easy-Spray column was maintained at 45℃ during the analysis. HCD analysis was performed on the Q Exactive instrument using a data-dependent acquisition method. The MS1 scans were acquired at 60000 resolution (200 m/z), automatic gain control (AGC) target of 1e6, and maximum ion injection time (IT) of 20 ms. Top 10 precursors were selected for MS/MS acquisition, with a resolution of 30000 (200 m/z), a 2 m/z isolation window, max IT of 100 ms, and AGC value of 5e4. Charge exclusion was applied to exclude unassigned and charge 1 species, and dynamic exclusion was used with a duration of 60 seconds.

### Proteomic data analysis

Protein identification and quantification were performed using Proteome Discoverer (version 3.1.0.638; Thermo Fisher Scientific) using the Rattus norvegicus fasta database (sp_incl_isoforms TaxID=10116; v2024-03-27), downloaded from UniProtKB and containing 9836 sequences, and a Contaminants fasta file with 379 common contaminant sequences. The data were processed using the common processing workflow for LFQ quantification and enhanced annotation for LFQ consensus workflow. The precursor mass tolerance was set to 10 ppm and the fragment mass tolerance to 0.02 Da with the enzyme set to Trypsin (full). A maximum of 2 missed cleavages was allowed. Dynamic modifications were set to oxidation (+15.995 Da) of methionine, N-terminal acetylation (+42.011 da), N-terminal methionine loss (−131.040 Da) and N-terminal methionine loss plus acetylation (−89.030 Da). Static modifications were set to carbamidomethylation (+57.021 Da) of cysteine. Percolator settings allowed a 1% false discovery rate (FDR) (strict) and 5% FDR (relaxed) for peptide and protein identifications. The time window for chromatographic peak alignment was set to 20 minutes, and normalization was based on the total peptide intensity of the samples. Quantification ratios were calculated using the SHR vehicle group as the denominator. Gene Ontology (GO) of biological processes was performed on differentially expressed proteins using Database for Annotation Visualization and Integrated Discovery (DAVID).

### General statistical analysis

Statistical analyses were conducted using GraphPad Prism 10. To assess statistical significance for BP and histology data, a one-way ANOVA was employed. Weekly BP data were analyzed using a repeated measures test. Myograph contractility data were analyzed using a non-linear regression model (log (agonist) vs. normalized response with a variable slope). Two-way ANOVA between two groups was used to compare difference between curves. Imaging data were evaluated using one-way ANOVA with a post hoc Tukey correction for multiple comparisons. For comparisons between two groups, a Student’s t-test was utilized. A p-value of less than 0.05 was considered statistically significant. Data are presented as mean ± SEM, with the number of samples (n) indicated in figure captions or as individual data points in the graphs.

---

**Supplementary Figure 1.**
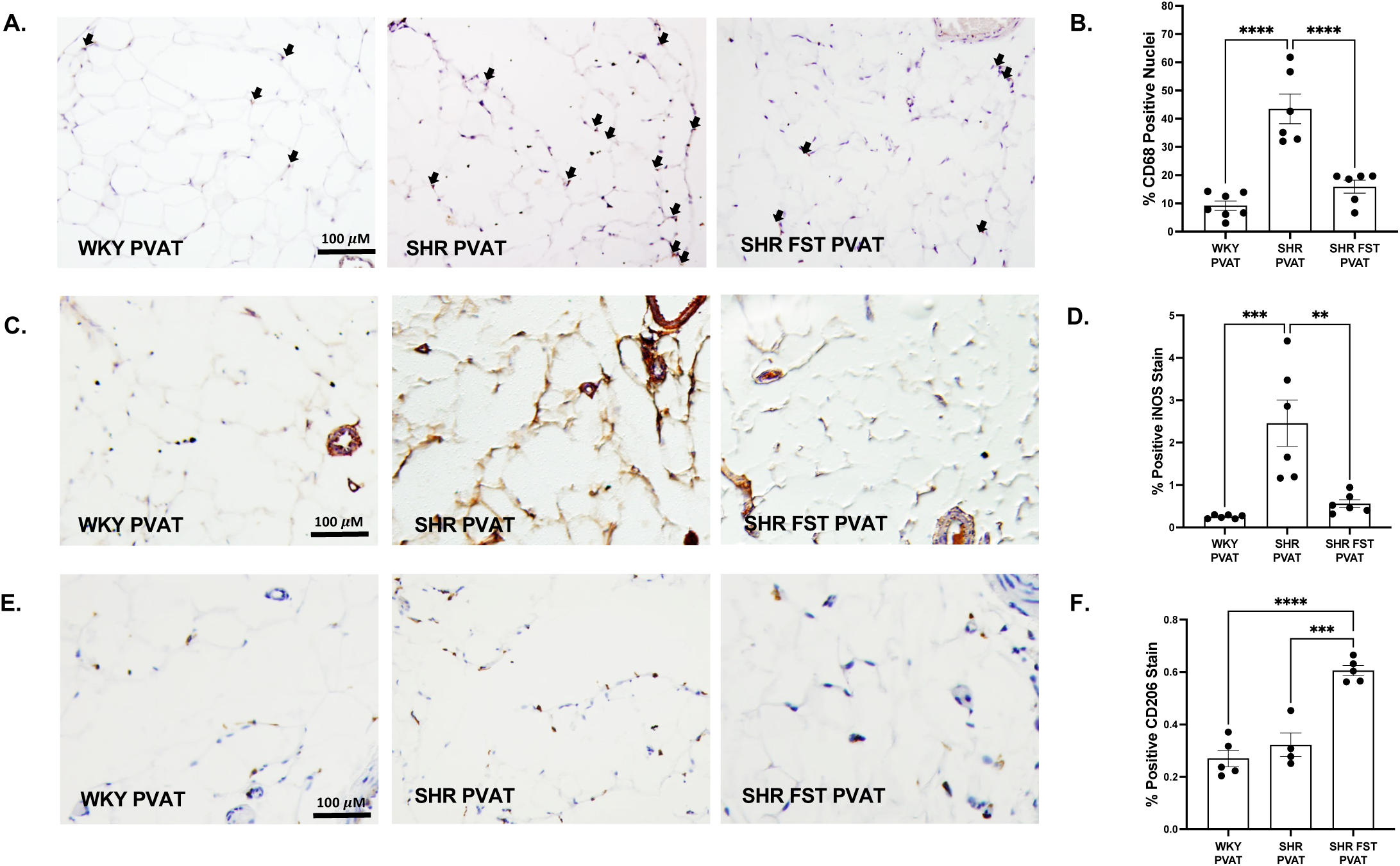
Follistatin inhibits PVAT inflammation in SHRs. Representative cross-sectional images of mesenteric PVAT from chronically treated rats stained for **A.** CD68, **C.** iNOS, and **E.** CD206, quantified in accompanying graphs **B., D., and F.** *\*\*p* <0.01, *\*\*\*p* <0.001, *\*\*\*\*p* <0.0001.

**Supplementary Figure 2.**
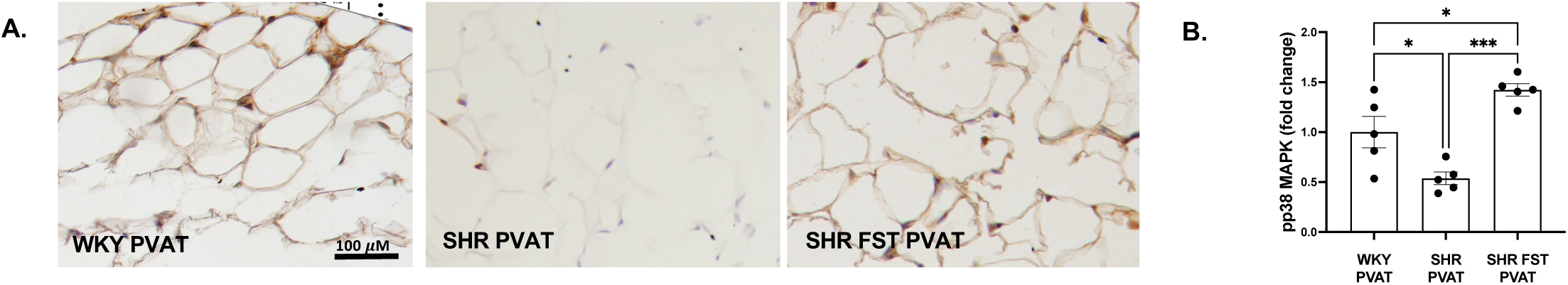
Follistatin increases phosphorylation of p38 MAPK. **A.** Representative cross-sectional images of mesenteric PVAT from chronically treated rats stained for phosphorylated p38 MAPK, quantified in **B.** *\*p*<0.05, *\*\*\*p* <0.001.

**Supplementary Figure 3.**
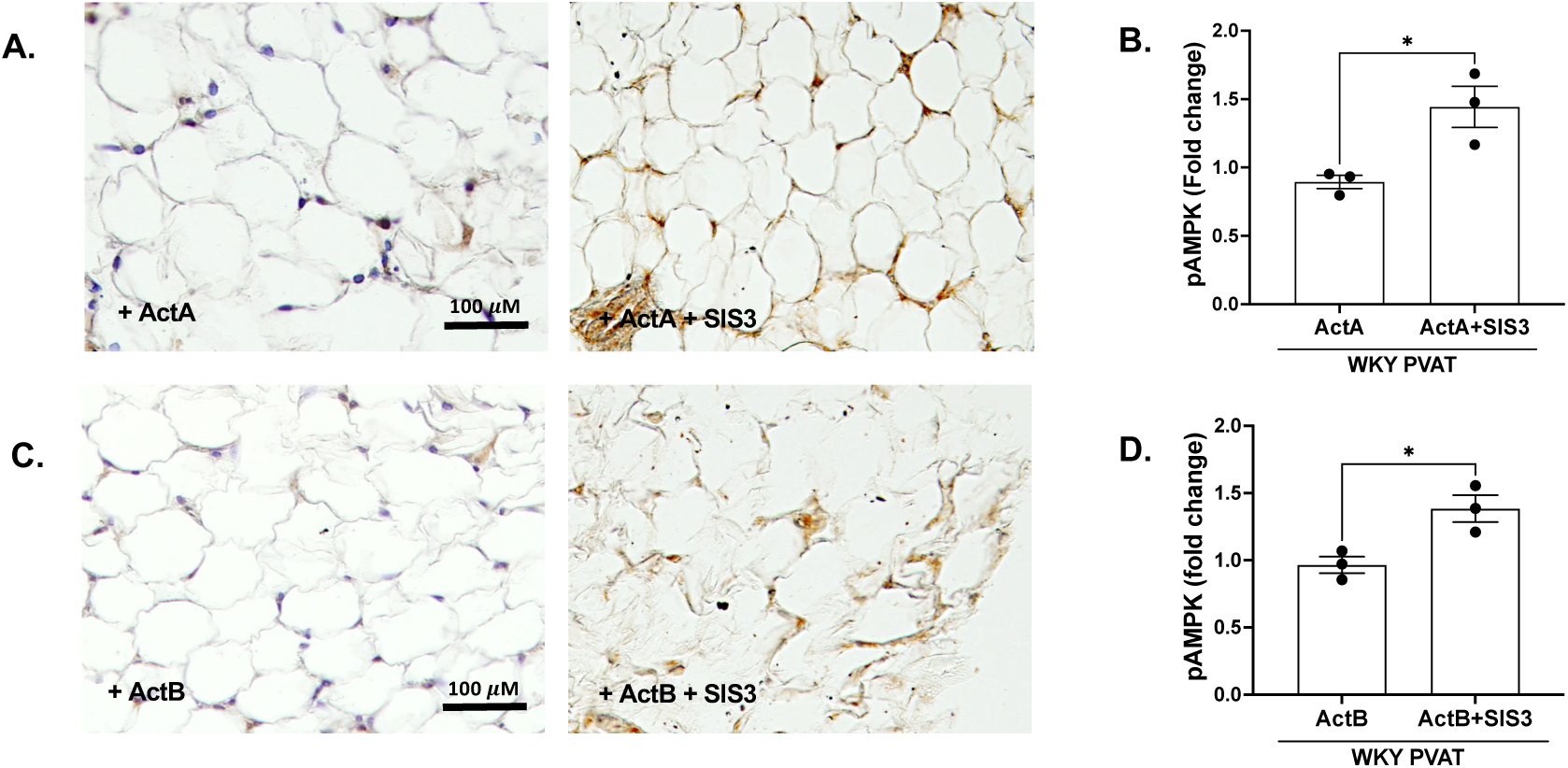
Smad3 inhibition increases AMPK phosphorylation. Representative cross-sectional images of WKY mesenteric PVAT treated ex vivo for 3 days with either **A.** activin A ± SIS3 and **C.** activin B ± SIS3 and stained for phosphorylated AMPK, quantified in **B.,** and **D.** *\*p*<0.05, *\*\*p* <0.01, *\*\*\*p* <0.001.

**Supplementary Figure 4.**
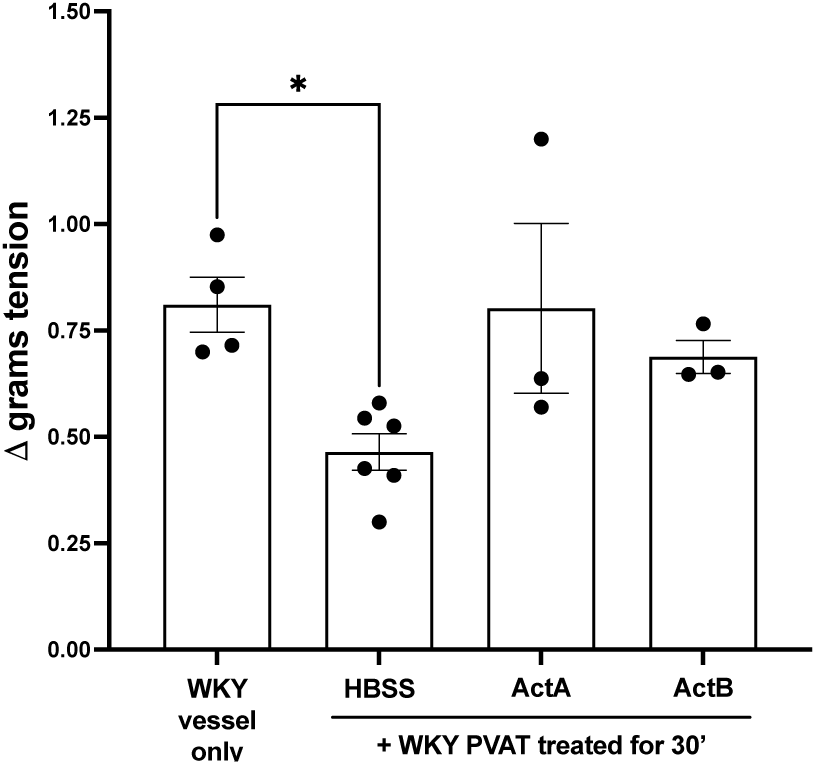
Activins reduce the anticontractile effect of WKY PVAT. Response to maximal KCl dose (125 mM) in WKY vessels incubated with detached mesenteric WKY PVAT pretreated with either HBSS, activin A or activin B for 30 minutes. Contractility values are from 1-2 rats per group, 1-3 vessels each rat. *\*p*<0.05.

**Supplementary Figure 5:**
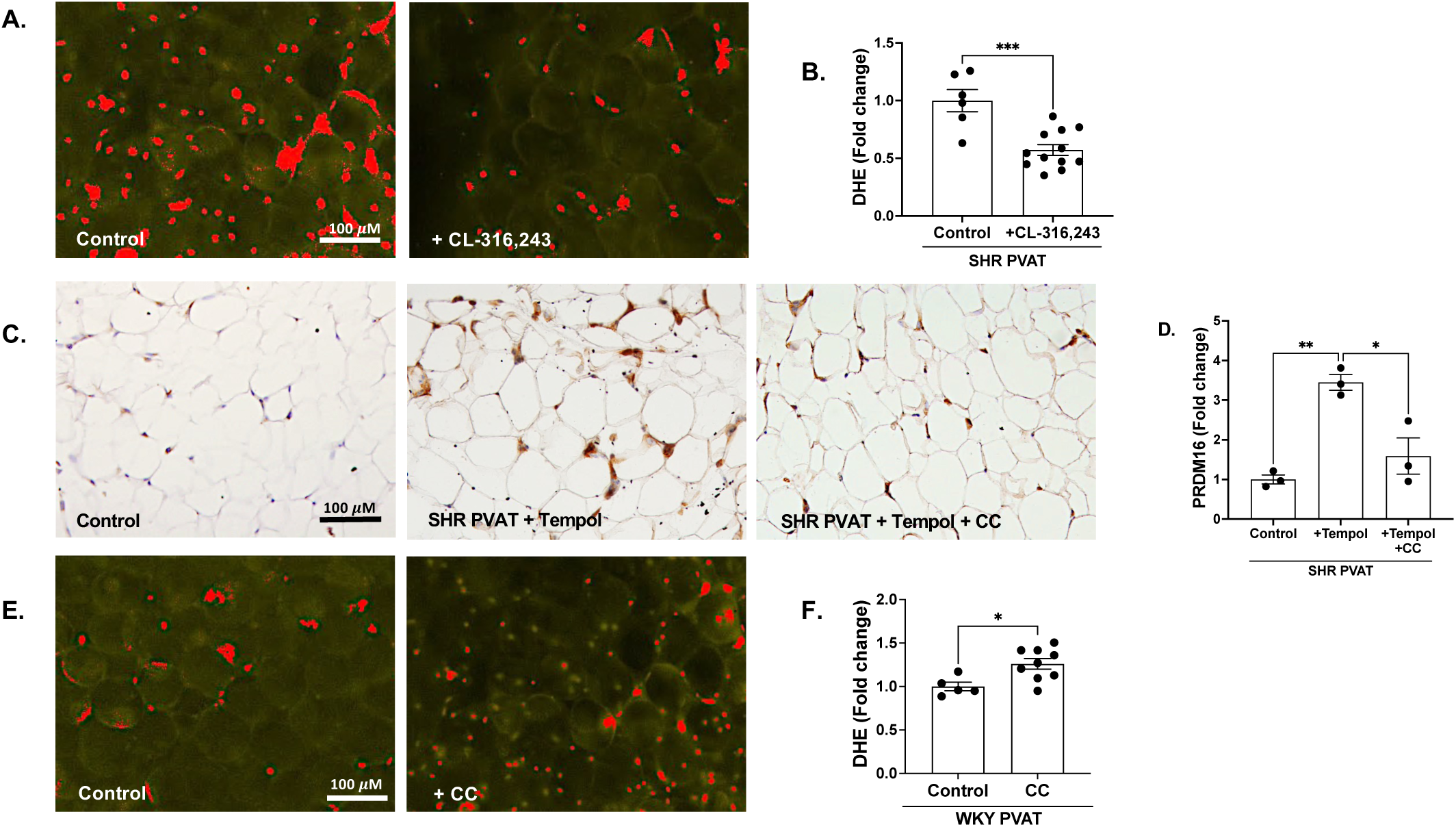
Relationship between oxidative stress, AMPK and browning. **A.** Representative fluorescence images of SHR mesenteric PVAT treated ex vivo for 3 days with either HBSS or CL-316,243 and stained with DHE, quantified in **B. C.** Representative cross-sectional images of SHR mesenteric PVAT treated ex vivo for 3 days with either HBSS, Tempol or Tempol with CC and stained for PRDM16, quantified in **D. E.** Representative fluorescence images of SHR mesenteric PVAT treated ex vivo for 3 days with HBSS or CC and stained with DHE, quantified in **F.** *\*p*<0.05, *\*\*p* <0.01, *\*\*\*p* <0.001.

**Supplementary Figure 6:**
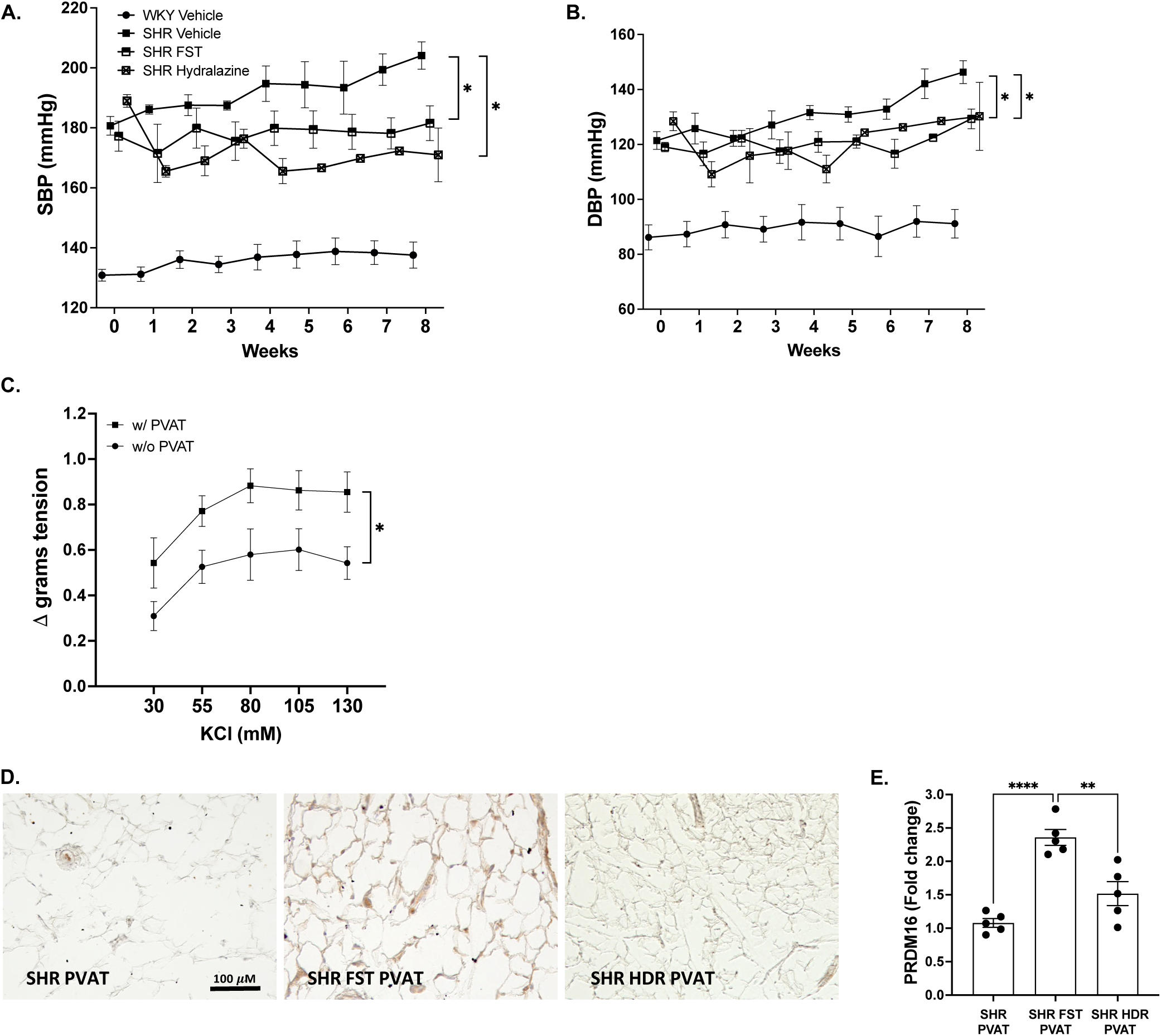
BP lowering does not improve vascular function or induce adipose tissue browning. Weekly **A.** systolic and **B.** diastolic BP measured by radiotelemetry in SHRs treated with hydralazine in drinking water for 8 weeks. WKY vehicle, SHR vehicle and SHR follistatin BP data are included from our previous treatment study for comparison. **C.** KCl dose response curves (30 – 130 mM) in first-order mesenteric vessels with or without PVAT from hydralazine-treated SHRs. **D.** Representative cross-sectional images of mesenteric PVAT from chronically treated SHRs stained for PRDM16, quantified in **E.** HDR, hydralazine; Contractility values are from 3-5 rats per group, 1-3 vessels each rat. *\*p*<0.05, *\*\*p* <0.01, *\*\*\*\*p* <0.0001.

**Supplementary Figure 7:**
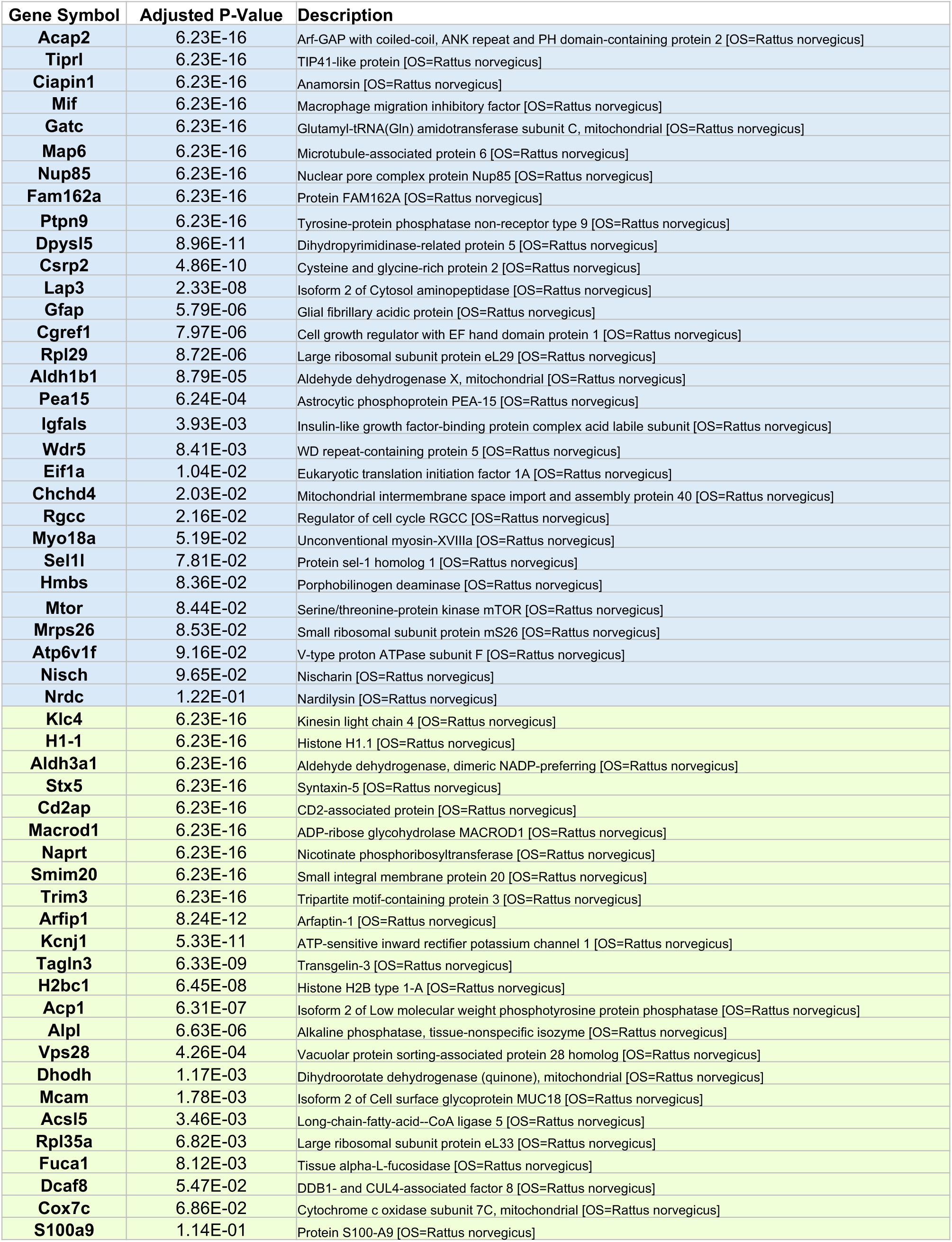
List of differentially expressed proteins regulated by follistatin. Adjusted p-values reflect changes in protein abundance in SHR FST PVAT vs. SHR PVAT, adjusted for multiple testing. Blue – upregulated proteins, green – downregulated proteins.

